# Recent Adversity Shapes Autonomic and Neural Threat Learning and Retrieval Under Imminent Threat: A Prospective Longitudinal Study

**DOI:** 10.64898/2026.07.21.739772

**Authors:** Maren Klingelhöfer-Jens, Manuel Kuhn, Tobias Sommer, Tina B. Lonsdorf

**Author notes:** Correspondence concerning this article should be addressed to Maren Klingelhöfer-Jens, Martinistrasse 52, 20246 Hamburg.

## Abstract

Recent adversity (RA) increases the risk of later mental health problems, but the mechanisms underlying this association remain largely unclear. Altered associative learning and specifically threat learning processes may constitute a potential pathway. Cross-sectional studies link RA to reduced threat-safety discrimination, a process crucial for adaptive functioning; however, longitudinal evidence allowing for more causal conclusions is lacking. In this prospective longitudinal study, 69 healthy participants completed fear acquisition, 24h-delayed extinction training and a reinstatement-test at two measurement time points six months apart (T0 and T1). We captured exposure to RA between T0 and T1 as well as autonomic and neural responses (SCRs and BOLD-fMRI), fear ratings, and salivary and hair cortisol at both assessments. During acquisition training, at the beginning of extinction (i.e., fear recall) and reinstatement-test at T1, RA-exposed individuals showed less autonomic threat-safety discrimination than unexposed individuals, mainly driven by blunted threat signal responding. These group differences predicted depression levels at the 1 and 1.5 year follow-ups and were mirrored by distinct activation in key fear-related brain regions, including striatal regions, thalamus, vmPFC, insula, and amygdala, whereas self-reported fear was unrelated to RA. In conclusion, across multiple outcomes, RA shapes threat learning and retrieval under conditions of (potential) imminent threat, with reduced autonomic threat discrimination prospectively predicting elevated depressive symptoms at 1- and 1.5-year follow-up. Overall, these findings implicate altered threat learning and retrieval processes as a potential mechanistic pathway through which RA becomes biologically embedded, potentially elevating psychopathological risk.

## Introduction

Recent adverse life experiences (recent adversity, RA) are among the most robust predictors of subsequent mental health difficulties - including the onset and aggravation of symptom severity as well as heightened vulnerability to relapse (1–5). Epidemiological evidence clearly implies that exposure to stressful or adverse events can drive transitions from subclinical to clinically relevant states by destabilizing remission (6,7). This underscores the urgent need to better understand the mechanisms through which exposure to recent adverse experiences shape vulnerability. A key candidate mechanism involves alterations in threat processing and associative learning (8,9). Adverse experiences may alter how threat-related cues are processed leading to changes in the expression and regulation of defensive threat responses and ultimately affective regulation and relapse risk. Addressing this question requires prospective and longitudinal experimental approaches that allow to track if and how threat-related associative learning changes as a function of exposure.

The fear conditioning paradigm (8) serves as a well-established translational model that targets core mechanisms of threat learning, expression and relapse. In differential fear conditioning protocols (8), a neutral stimulus is repeatedly paired with an aversive unconditioned stimulus (US; e.g., an electrotactile shock), thereby becoming a conditioned stimulus (CS+) and establishing a CS+/US association through which the threat signal gains the power to predict the US (acquisition training). In contrast, a second stimulus, the CS−, is never paired with the US and may be conceptualized as safety signal. Acquisition training is considered an experimental model for the acquisition of fear (10). During extinction training, the conditioned stimuli (CS+ and CS-) are presented without the US, leading to a gradual reduction of the conditioned response. Importantly, extinction training - considered as a key mechanism underlying exposure-based treatments (10–13) - does not erase the fear memory (CS+/US association) but is considered to create a competing inhibitory extinction memory trace (CS+/no US association, 10,14). Extinction can be followed by a reinstatement phase (15) during which the re-presentation of the US can trigger a return of fear (ROF). Reinstatement has been suggested to serve as a laboratory model for clinical relapse (13,16). During the reinstatement-test phase conditioned responses are re-assessed to distinguish between extinction retention (i.e., the absence of conditioned responses) or the ROF (8,17).

Initial evidence from cross-sectional work suggests that exposure to RA is associated with reduced threat-safety discrimination (i.e., CS+ > CS-) during fear acquisition training (18) as well as during fear recall (i.e., early extinction trials in a delayed extinction protocol) and reinstatement-test (16). Furthermore, more pronounced exposure to stressful events was associated with a dose-related increase in neural differentiation between threat and safety cues in the amygdala during acquisition training (18), and linked to reduced neural engagement during fear recall and reinstatement across key regions involved in threat processing and regulation - including the amygdala, hippocampus, and thalamus (16). Of note, in both studies, reduced discrimination between threat and safety was driven by blunted responding to the anticipatory threat signal (CS+). Previous work suggests that threat-safety discrimination abilities are linked to healthy functioning and resilience (19,20). Thus, reduced discrimination may represent an important mechanism linking RA to mental health symptoms, with altered threat learning processes providing a plausible pathway through which adversity may become biologically embedded.

Critically, however, evidence for effects of RA on threat learning mechanisms originates entirely from cross-sectional work which precludes inferences about directionality and leaves open whether observed effects reflect potentially pre-existing group differences or indeed consequences of exposure to adversity. To fill this gap, we conducted a prospective longitudinal study using a differential fear conditioning paradigm in a large sample of mentally healthy adults. We used an identical procedure (day 1: acquisition training; day 2: extinction training, reinstatement and reinstatement-test) at two measurement time points (T0, T1) six months apart and combined autonomic (SCRs) and subjective (fear ratings) indices with BOLD fMRI to capture the neural mechanisms through which RA may become biologically embedded in threat processing systems. Further, we tested if threat-safety discrimination at T1 predicts anxiety and depression levels at a 1-, 1.5-, and 2-year follow-up.

Based on cross-sectional results, we hypothesize that individuals exposed to RA in the six months interval will show diminished threat-safety discrimination at T1 (e.g., after exposure, controlled for T0 responding) - specifically during acquisition training, at fear recall as well as during reinstatement-test. Moreover, we hypothesize that threat-safety discrimination predicts higher anxiety and depression levels across the follow-up period. To further characterize potential group differences at the endocrine level, we assessed salivary and hair cortisol as markers of acute stress reactivity and longer-term hypothalamic–pituitary–adrenal axis activity, respectively, expecting lower salivary cortisol reactivity during the experiment (21), but higher hair cortisol levels (22) in individuals exposed to RA as compared to unexposed individuals.

## Methods and Materials

### Participants and Experimental Design

69 healthy participants (female_M,T1_ = 40, male_M,T1_ = 29, age_M,T1_ = 25.30, age_SD,T1_ = 3.83, age_range,T1_ = 19 - 33) screened for the absence of adversity during childhood and/or adolescence were selected from a large recruitment cohort (23). More detailed information on sample characteristics and participant exclusions is provided in the Supplement (see Supplementary Figure 1 and Supplementary Table 1). Participants underwent a differential fear conditioning paradigm similar to (16) using fractals as CSs and an electrotactile stimulation as US. The experimental phases included habituation and acquisition training on day 1, followed by a 24-h delayed extinction phase and a reinstatement-test on day 2 (baseline, T0). The identical procedure was repeated after 6 months (T1). Details on the acquisition of SCRs, BOLD fMRI, fear ratings, salivary and hair cortisol are described in the supplement. Furthermore, 1-, 1.5-, and 2-year after T1, participants completed the State-Trait Anxiety Inventory (STAI-T, 24) and Beck Depression Inventory-II (BDI-II, 25) at online follow-up assessments (T2-T4) to capture anxiety and depression levels. All participants gave written informed consent to the protocol which was approved by the local ethics committee (PV 5157, Ethics Committee of the General Medical Council Hamburg, Germany).

### Quantification of recent adversity

Participants were classified as exposed (RA) or unexposed (no RA) to at least one negative life event in the 6 months time interval between T0 and T1 based on the List of Threatening Experiences (LTE, 26), which was administered after the experimental session at T1 on day 2.

### Data Analyses of Ratings, Psychophysiology, and Salivary Cortisol

For SCR and fear ratings, linear mixed-effects models were fitted separately for acquisition training, extinction training, fear recall and reinstatement-test, entering CS type and RA as fixed effects, along with their interactions. The main analyses were conducted using the lmer() function from the lme4 package (27), specifying random intercepts for participants. Fixed effects included baseline responses (T0) which were centered within individuals as a covariate, which allowed controlling for initial baseline differences. Predictions of anxiety and depression levels were tested by using linear regression models with differential SCRs and extracted neural parameter estimates from the CS+ versus CS− contrast for acquisition, fear recall, and reinstatement as predictors and STAI-T/BDI-II scores at T2 - T4 as outcomes.

For salivary cortisol, the area under the curve with respect to increase (AUCi) was calculated according to (28) using the trapezoidal rule, subtracting the first measurement as baseline. Absolute cortisol levels across time-points are reported in the Supplement (see Supplementary Figure 2). A linear model with AUCi at T1 as the dependent variable, RA as the independent variable, and AUCi at T0 as a covariate was used.

## Results

Unless otherwise stated, all reported statistical results are based on analyses of T1 outcomes while controlling for baseline (T0) values - with the exception of manipulation checks, which were analyzed separately for T0 and T1.

### Manipulation check

Across outcome measures and time-points (T0 and T1), successful threat acquisition was confirmed at the group level by significant CS discrimination (i.e., CS+ > CS-) and successful extinction by a significant reduction of CS discrimination (SCRs: from early to late extinction; fear ratings: pre- to post-extinction). Yet, extinction was incomplete as CS discrimination in fear ratings was - despite this reduction - still significant at the end of extinction training (see Figure 2 and Supplementary Table 3). Neural activation to the CS+ was higher as compared to the CS- during acquisition training in expected brain regions typically linked to conditioned discrimination (see CS+ > CS- contrast T-maps on NeuroVault; https://identifiers.org/neurovault.collection:24143).

### Salivary and hair cortisol

Participants exposed to RA between T0 and T1 showed a significantly flatter salivary cortisol decline (AUCi) over the course of the experiment on day 1 as compared to unexposed participants (i.e., acquisition training, Figure 1A, *t*(55) = 2.20, *p* = .032), but not on day 2 (i.e., extinction training and reinstatement-test, Figure 1B, *t*(53) = 0.39, *p* = .695). Changes of cortisol (AUCi) were unrelated between both assessment time points (T0, T1) for both experimental days (day 1: *t*(55) = 1.12, *p* = .269, day 2: *t*(53) = 1.02, *p* = .314), supporting the interpretation that adversity-related differences in acute endocrine stress reactivity emerged between T0 and T1 rather than reflecting RA-unrelated group differences. Analyses of absolute cortisol levels and hair cortisol levels (see Figure 1C) are reported in the Supplement (see Supplementary Figure 2 and Supplementary Table 2). In brief, no significant group differences were observed in hair cortisol levels (*F*(1,68.42) = 0.80, *p* = .374), despite descriptively somewhat higher levels in exposed individuals.

**Figure 1:**
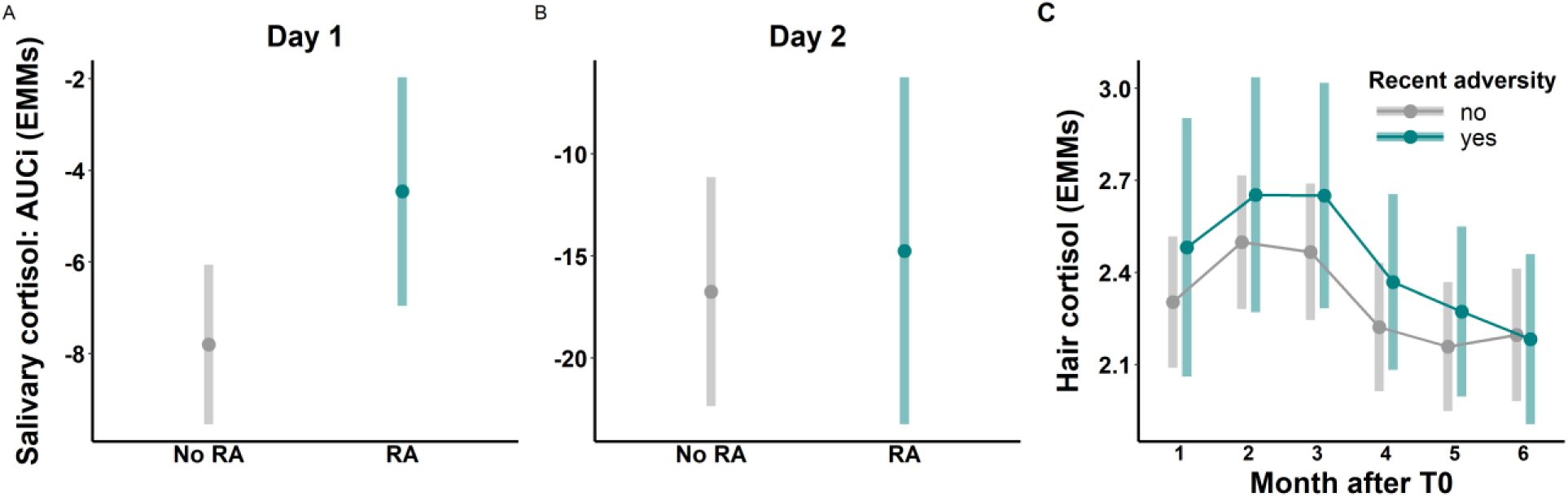
Salivary cortisol responses at T1, quantified as area under the curve with respect to increase (AUCi) and adjusted for baseline levels (T0), are shown for (A) day 1 and (B) day 2, reflecting changes from baseline across the experimental sessions (estimated marginal means, EMMs; see methods for description of the statistical models). Higher values indicate a flatter decline in cortisol over time. On day 1, three saliva samples were taken with one sample before attachment of the electrodes (sample 1) and two samples after acquisition training (sample 2: 20 min after sample 1; sample 3: 30 min after sample 1). On day 2, five saliva samples were extracted: before attachment of the electrodes (sample 1), after RI-test (sample 2: 30 min after sample 1), subsequent to structural scans (sample 3: 50 min after sample 1), and as well as 60 min and 70 min after sample 1 (i.e., sample 4 and 5). (C) EMMs of hair cortisol concentrations across different hair segments. Each of the six segments represents one month within the six-month period between T0 and T1.

**Figure 2:**
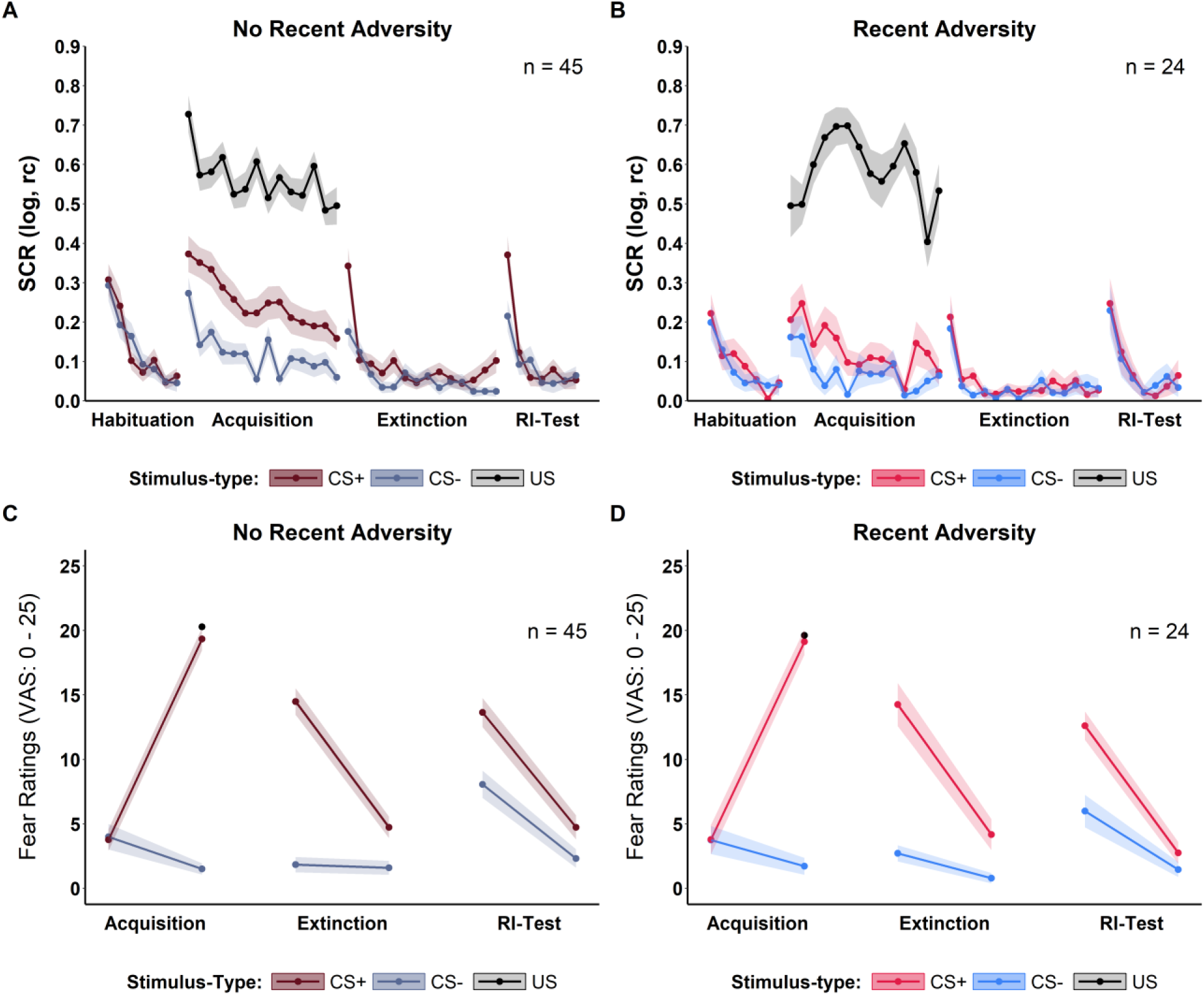
Trial-by-trial SCR data (A and B) to the CS+ (in red), CS- (in blue) and US (in black) and pre- as well as post-phase fear ratings (C and D) at T1 for the exposed (left) and unexposed (right) group. Shaded ribbons represent 95% CIs. CS = conditioned stimulus, US = unconditioned stimulus.

### Effects of RA exposure on SCRs, BOLD fMRI and fear ratings

Individuals exposed to RA showed significantly reduced CS discrimination in autonomic responding during acquisition training as compared to unexposed individuals (CS type x RA interaction *F*(1,66) = 4.72, *p* = .033; exposed: *t*(66) = 1.19, *p* = .240; unexposed: *t*(66) = 3.58, *p* < .001 at T1 (see Figure 2, Supplementary Figure 3 and Supplementary Table 4). This was driven by significantly attenuated average SCRs to the anticipatory threat cue (i.e., CS+) in the exposed group (*t*(88.98) = 2.91, *p* = .005), while SCRs to the safety cue (i.e., CS-) were not significantly different between both groups (*t*(88.98) = 1.26, *p* = .211). Crucially, despite these differences in anticipatory threat responding to the CS+, no group differences in the reactivity to the unconditioned threat cue (i.e., SCRs to the US; *t*(67) = -0.66, *p* = 0.514, see also Figure 2) or US aversiveness ratings were observed (see Supplementary Table 1). This pattern indicates that the aversive learning signal itself (i.e., US) was perceived and processed comparably in both groups, implying that group differences are restricted to and emerge only at the level of anticipatory associative threat learning.

The significant group differences in autonomic CS discrimination during acquisition training were reflected in corresponding group differences in CS discrimination in neural activity in areas implicated in threat processing including striatal areas (left and right nucleus accumbens, left putamen, right pallidum), left thalamus, and the ventromedial prefrontal cortex (vmPFC, see Table 1A and Figure 3A). Yet, while CS discrimination in autonomic responding was reduced in exposed as compared with unexposed individuals, CS discrimination in neural activity was enhanced (see Table 1A and Figure 3A). Extracted parameter estimates indicate that this might be driven by both, enhanced responses to the anticipatory threat (i.e., CS+) and reduced responses to the anticipatory safety signal (i.e., CS-)

**Figure 3:**
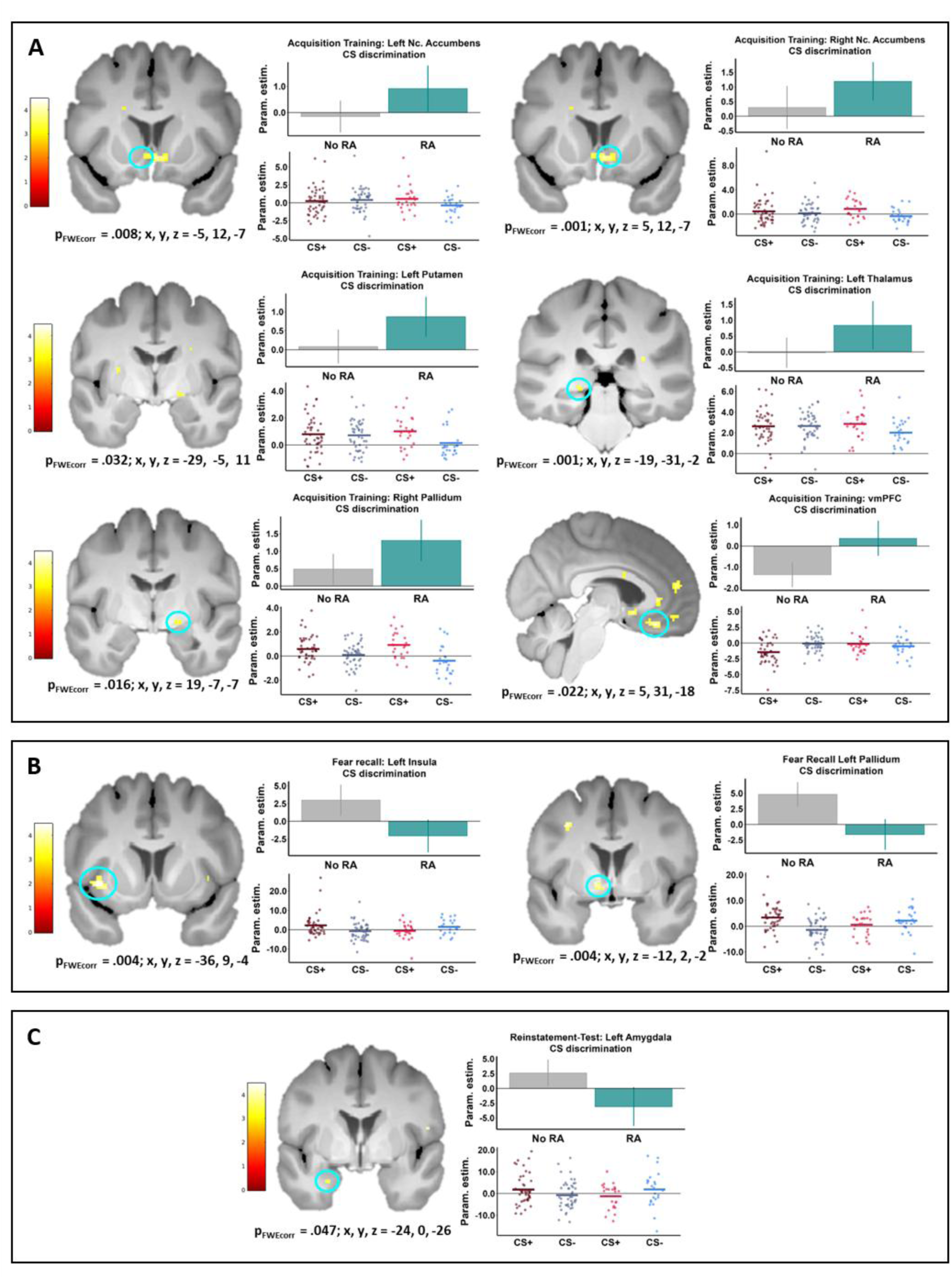
Neural activation at T1 reflecting group differences in CS discrimination during acquisition training (A), fear recall (B), and reinstatement (C) on a visualization threshold of p < 0.001 and corresponding beta values for CS discrimination (upper panel) and CS+ and CS- separately (lower panel). For acquisition training, significant effects were observed for the contrast that tested for higher CS discrimination in the exposed versus unexposed group at T1 relative to T0, while for fear recall and reinstatement, the reverse pattern was observed. Error bars represent 95% CIs (standard error of the mean). CS = conditioned stimulus.

**Table 1.**
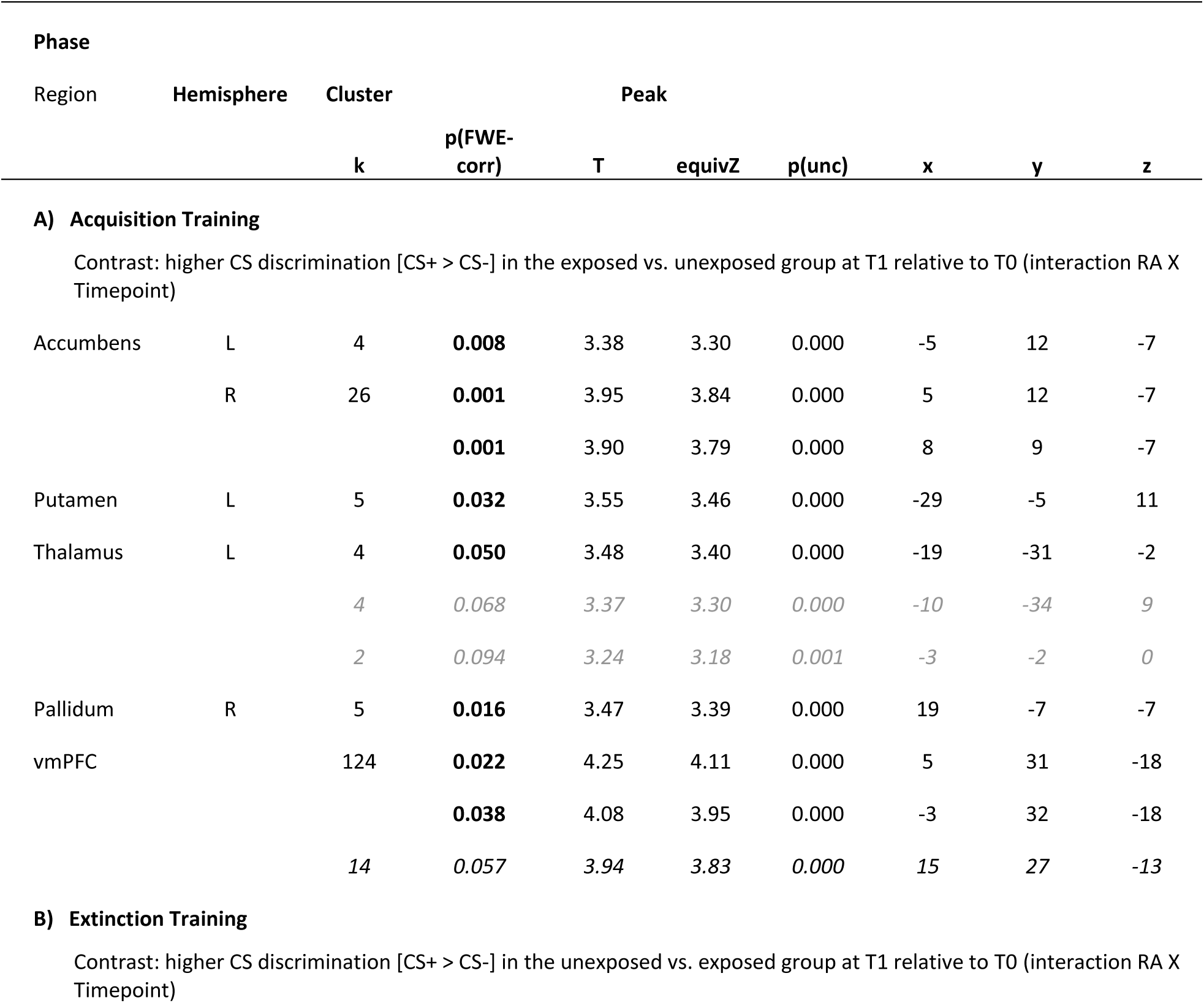
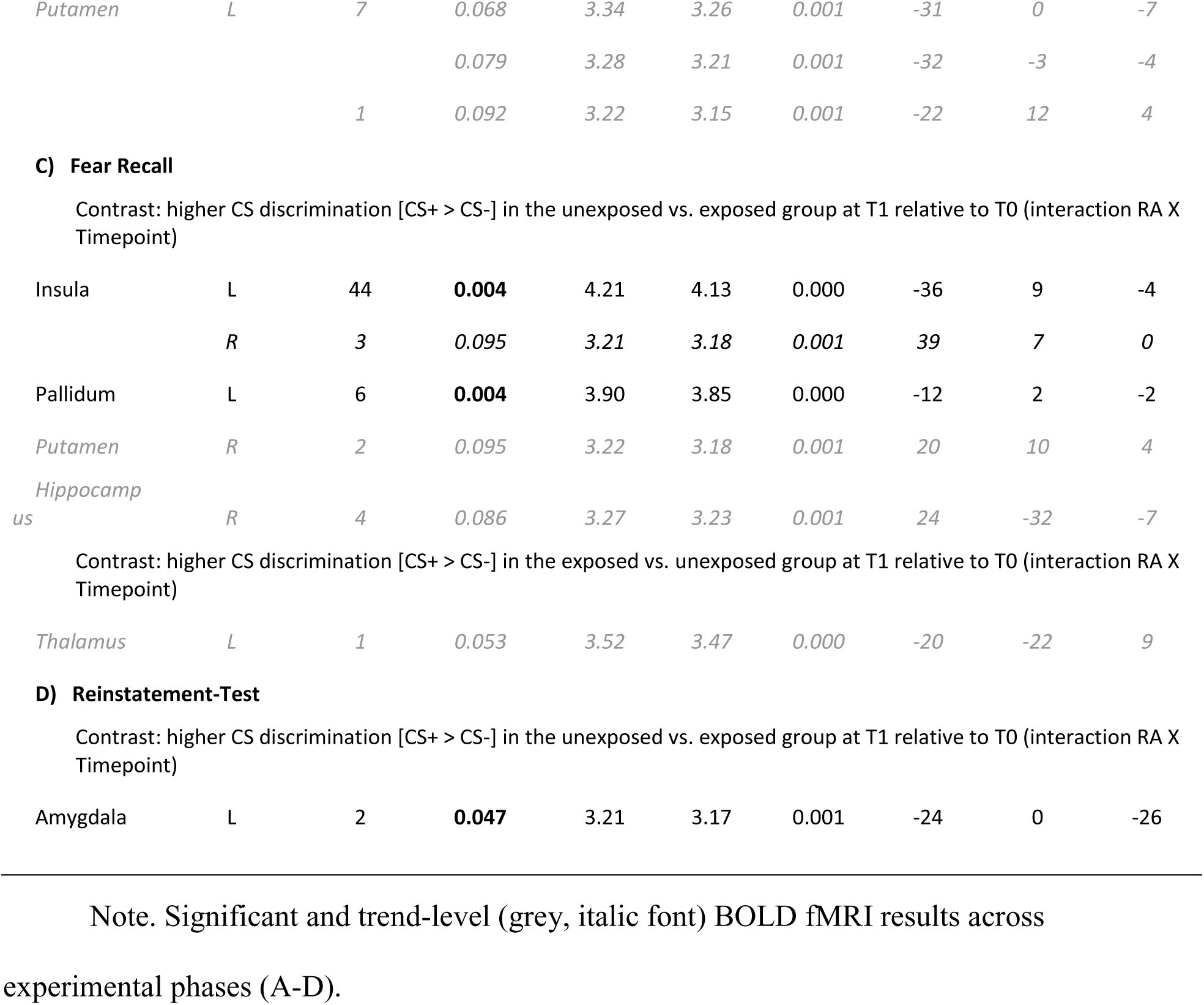
BOLD fMRI results.

As during acquisition training, exposed individuals showed less autonomic CS discrimination as compared to unexposed individuals during both fear recall (i.e, first extinction trial of 24h-delayed extinction) and reinstatement-test (see Figure 2, Supplementary Figure 3 and Supplementary Table 4); i.e, CS type x RA; fear recall: *F*(1,131.66) = 7.40, *p* = .007; reinstatement-test: *F*(1,200.25) = 5.13, *p* = .025). As during acquisition training, this was driven by significantly attenuated autonomic responding to the conditioned threat cue (i.e., CS+) in the exposed group (fear recall: *t*(101.03) = 2.35, *p* = .020; reinstatement-test: *t*(130.22) = 2.34, *p* = .021 in absence of significant group differences in responding to the safety signal (fear recall: *t*(101.13) = −0.09, *p* = .926; reinstatement: *t*(130.22) = −0.22, *p* = .828.

Also as during acquisition training, significant group differences were mirrored in several brain regions known to be implicated in threat processing including the left insula, left pallidum (fear recall, see Table 1C and Figure 3B), and left amygdala (reinstatement-test; see Table 1D and Figure 3C)) and at non-significant trend-level in the left putamen, hippocampus and right thalamus (fear recall, see Table 1C). Yet, in contrast to the dissociation between autonomic and neural CS discrimination observed during acquisition, group differences in CS-related neural activation during fear recall and reinstatement were reduced (with the exception of the thalamus at fear recall) in the exposed group and aligned with the autonomic pattern, with both measures showing the same direction of effects.

Of note, reduced autonomic, but not neural threat-safety discrimination at T1 prospectively predicted higher depression levels (BDI-II) at 1 year (fear recall; *t*(63) = −2.07, *p* = .043 and 1.5 year (fear acquisition; *t*(62) = −2.10, *p* = .039) follow-up (see Figure 4). However, this predictive association was no longer significant at the 2-year follow-up (see Supplementary Table 5 and 6 for details). No significant predictions were observed for anxiety symptoms, as assessed by the STAI-T.

In contrast to autonomic and neural levels, no significant group differences were observed on any experimental day for fear ratings across experimental phases (see Figure 2, Supplementary Figure 4 and Supplementary Table 7) and for extinction training across outcomes measures (SCRs: Figure 2, Supplementary Figure 3 and Supplementary Table 4; BOLD fMRI: Table 1 and Figure 3).

(ref:figure-caption-combined-scatter-clinical-prediction) Significant and trend-level (grey, italic font) BOLD fMRI results across experimental phases (A-D).

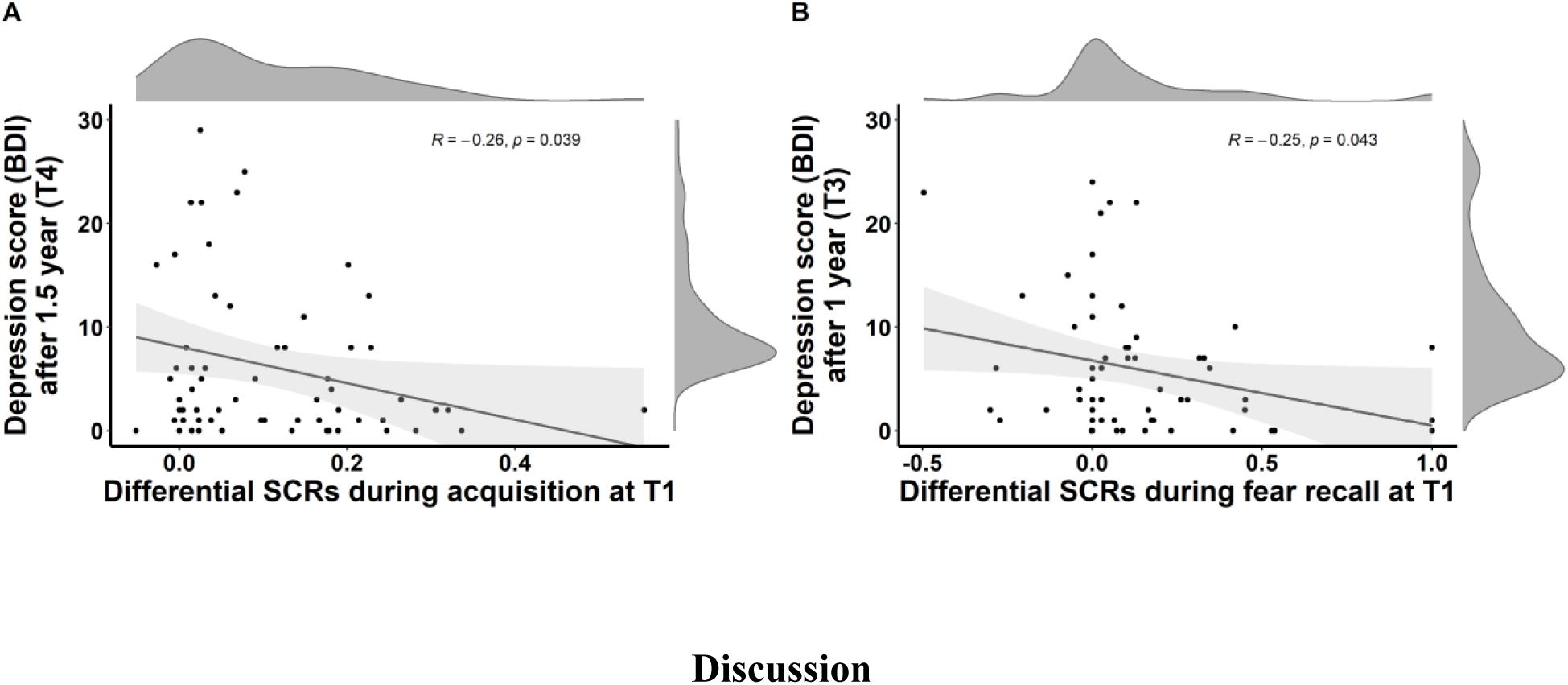

## Discussion

This prospective longitudinal study investigated how recent adversity (RA) shapes threat learning across acquisition, extinction, and reinstatement as a mechanistic model of adversity-related vulnerability. RA was followed by reduced threat-safety discrimination in autonomic and neural responses during experimental phases characterized by (potentially) imminent threat (i.e., acquisition, 24h-delayed fear recall, and reinstatement), which, in turn, predicted depressive symptoms at the 1- and 1.5-year follow-ups. This pattern of reduced threat-safety discrimination was driven by blunted anticipatory responses to the conditioned threat cue (CS+), while safety-related responding (CS-) remained intact, corroborating previous cross-sectional work (16,18). Importantly, RA did not alter autonomic responses to the unconditioned threat (US), indicating preserved defensive threat reactivity. Together with the absence of RA effects on explicit subjective fear ratings, this suggests that RA specifically affects implicit anticipatory threat responding.

Safety learning (i.e., extinction training), however, appeared to be unaffected by RA, as indicated by the absence of RA effects. Combined with the re-emergence of group differences during fear recall and reinstatement, these findings suggest that effects of RA are not broadly distributed across all fear-related mechanisms but emerge specifically under conditions of imminent threat - potentially reflecting persistent differences in threat - but not the extinction - memory trace representation and expression (14). A striking exception to the general pattern of reduced threat-safety discrimination is a relatively higher neural threat-safety discrimination in individuals exposed to RA only during acquisition training, resulting in a remarkable dissociation between peripheral physiological arousal and central neural processing in areas implicated in threat processing (29) - including the thalamus, striatal areas (Ncl. accumbens, putamen, pallidum) and the vmPFC. Interestingly, a similar dissociation during threat learning was reported in cross-sectional work, with RA-exposed individuals showing reduced differential fear ratings alongside enhanced neural discrimination in the left amygdala (18).

The enhanced differential neural activation during acquisition training observed in our work may indicate that RA exposure biases early processing toward anticipatory threat cues. The thalamus may support enhanced detection and prioritization of threat-relevant inputs (30,31), while striatal regions may amplify threat cue signaling through outcome probability computations (32–34). Such preferential threat processing may represent a mechanism through which RA becomes biologically embedded. Importantly, despite this enhanced threat–safety discrimination at a neural level, downstream autonomic anticipatory threat responses (CS discrimination and SCRs to the CS+) were attenuated following RA exposure. This attenuation may reflect enhanced vmPFC-mediated inhibition of fear expression (35–37), consistent with the increased anticipatory threat cue–evoked vmPFC activation observed in individuals exposed to RA in our work. Thus, taken together, the dissociation of neural activation and autonomic responding during acquisition training might indicate a decoupling between more cognitive (thalamic) salience- and (striatal) evaluation-related threat-safety discrimination and physiological output following RA.

In the short term, such a dissociation may be adaptive but could entail substantial long-term costs. In environments characterized by adversity, the reliable and early detection and discrimination of threat-predictive cues may be particularly important. This is consistent with our observation of enhanced neural discrimination within salience- and evaluation-related regions, potentially indicative of a prioritization of threat over safety learning. At the same time, reduced autonomic mobilization in anticipation of potential threat may be metabolically economical in order to preserve physiological resources for responses to imminent threat. In line with this interpretation, autonomic responding to actual threats (i.e., the US) remained intact in our study. Together, such decoupling may reflect an adaptive short-term strategy with potential long-term costs, as persistent divergence between cognitive and autonomic threat signal processing may compromise interoceptive coherence and ultimately emotion regulation. Supporting this possibility, individuals exposed to RA showed reduced insular threat–safety discrimination during fear recall. Given the role of the insula in integrating bodily and threat-related signals (38), altered insular threat–safety discrimination may indeed reflect disrupted brain–body integration following RA. Although direct evidence linking RA to interoceptive dysfunction is currently lacking, findings from chronic stress (e.g., 39,40,41) as well as exposure to early adversity (42) research suggest that adversity may disrupt interoceptive processes. In addition, impaired interoception has been also implicated across multiple psychiatric conditions, including anxiety, depression, schizophrenia, and PTSD (for overviews, see 43,44). Thus, while speculative, our findings highlight a potential pathway linking RA to psychopathology.

Strikingly, the above described dissociation between autonomic and neural responses was confined during threat learning. During fear retrieval 24h later (i.e., fear recall) and reinstatement, however, adversity-exposed individuals showed convergent blunting in threat-safety discrimination across both levels of analysis. More precisely, reduced differential activity in the pallidum and insula (during fear recall) as well as amygdala (during reinstatement) may reflect altered processing of conditioned threat relative to safety cue. This pattern is consistent with the contribution of these regions to value-related processing, threat evaluation, and autonomic aspects of fear expression (45–50). Of note, our results suggest that the pallidum may play a phase-dependent role in threat processing, in line with evidence that basal ganglia involvement varies across contexts and task demand (51,52). Here, pallidal threat-related responses were enhanced during threat learning but reduced during fear retrieval, suggesting dynamic modulation of threat processing across memory phases.

Interestingly, the blunted CS+ responding observed during threat learning and retrieval following RA closely mirrors patterns previously reported in individuals exposed to childhood adversity (CA) (23,53–55). First, in both CA and RA, adversity-related alterations appear restricted to phases involving imminent threat, such as acquisition and generalization (23,56–59), while extinction (i.e., safety) learning is preserved. Second, at a neural level, CA and RA-related differences in threat processing manifest most pronouncedly in striatal regions (56). This converging evidence may suggest that blunted anticipatory responding to threat cues may represent a broader signature of adversity exposure rather than being specific to CA or RA. However, this distinction remains unresolved to date, as many studies focusing on CA include samples of young adults (53) - as in this work - for which recent exposure to adversity may fall within a developmental period that would at the same time classify as late adolescents.

Of note, reduced autonomic threat-safety discrimination during threat learning and recall predicted higher depressive - but not anxiety - symptoms at 1- and 1.5-year follow-ups, suggesting threat–safety discrimination as a potential marker of vulnerability. This aligns with prior work linking intact threat–safety discrimination to healthy functioning and resilience (19,20). However, the absence of this association at 2 years follow-up may indicate that these effects are time-limited and thus potentially reversible.

Despite the strengths of this longitudinal design, which allowed us to examine prospective associations over time, some aspects require further investigation. A key finding of our study is that RA modulates fear retrieval processes (i.e., fear recall and reinstatement). However, it remains unclear whether these RA-related effects reflect truly retrieval-specific alterations or carry-over effects from RA-related differences during threat acquisition, or both. An interesting avenue for future studies might be to disentangle these mechanisms by tightly controlling threat learning processes, for example through explicit CS–US contingency instructions. Further, it remains unclear whether the observed neural differences extend also to explicit measures of contingency learning. Future studies incorporating trial-by-trial CS–US contingency ratings could clarify whether neural markers of threat discrimination are mirrored specifically in anticipatory autonomous responding or also in conscious threat learning.

In conclusion, we provide compelling evidence from a prospective longitudinal study covering multiple levels of analysis that suggests that under imminent threat, RA selectively shapes anticipatory threat responding, its retrieval and reinstatement following 24h consolidation. Furthermore, reduced threat discrimination at an autonomic level prospectively predicted elevated depressive symptoms at a 1 and 1-5 year follow up. Overall, our findings identify threat learning and retrieval processes as a potential mechanism through which RA becomes biologically embedded, thereby increasing vulnerability to psychopathology.

## Acknowledgements

The authors would like to thank Claudia Immisch, Janne Nold, Kevin Rozario, and Habiba Schiller for help with data collection, Karoline Rosenkranz for help with data preprocessing, and Mareike Clos for help with BOLD fMRI data analysis.

## Author contributions

The authors made the following contributions. Maren Klingelhöfer-Jens: Conceptualization, Data curation, Software, Formal analysis, Visualization, Methodology, Writing - Original Draft Preparation; Manuel Kuhn: Data curation, Software, Investigation, Methodology, Writing - Review & Editing; Tobias Sommer: Methodology, Formal analysis, Writing - Review & Editing; Tina B. Lonsdorf: Conzeptualization, Resources, Supervision, Funding acquisition, Methodology, Writing - Original Draft Preparation.

## Funding

Deutsche Forschungsgemeinschaft (INST 211/633-2): Tina B. Lonsdorf

Deutsche Forschungsgemeinschaft (LO 1980/4-1): Tina B. Lonsdorf

## Previously published data

SCR and rating data have been reported in a previous methodological study investigating the longitudinal reliability and cross-phase predictability of these measures within our fear-conditioning paradigm (60).

## Use of AI technologies

All content was written by the authors. UHHGPT (the AI service of the University of Hamburg) and ChatGPT were used solely for language refinement and grammar checking of the manuscript. All AI-assisted edits were reviewed and approved by the authors.

## Data availability

The data that support the findings of this study and the R Markdown files that generate this manuscript are openly available in Zenodo at 10.5281/zenodo.21472062.

