## Supplementary Material for "Recent Adversity Shapes Autonomic and Neural Threat Learning and Retrieval Under Imminent Threat: A Prospective Longitudinal Study"

### Appendix

#### Methods

##### Participants

In total, 120 participants ($female_{M,T0}$ = 79, $male_{M,T0}$ = 79 and 41, $age_{M,T0}$ = 24.68, $age_{SD,T0}$ = 3.70, $age_{range,T0}$ = 18 - 34) were a priori selected from a large participant pool (e.g., 23, using the Hamburg sample only) recruited to provide well-characterized participants for follow-up studies in the context of the Collaborative Research Center SFB TRR 58. For the present study, the selection was based on the absence of traumatic experiences during childhood as assessed by the Childhood Trauma Questionnaire (1,2): Hence, only participants with emotional abuse < 13, physical abuse < 10, sexual abuse < 8, emotional neglect < 15, and physical neglect < 10 were included (1,3). Exclusion criteria were age under 18 or over 50, regular medical (except oral contraceptives) or illegal substance intake, chronic diseases, and neurological/psychiatric disorders. Participants were right handed and had normal or corrected to normal vision. For sample characteristics of the sample included here, see Supplementary Table 1. All participants gave written informed consent to the protocol which was approved by the local ethics committee (PV 5157, Ethics Committee of the General Medical Council Hamburg, Germany). The study adhered to the principles outlined in the Declaration of Helsinki. All participants were unfamiliar with the experimental setup prior to the study. In total, the study involved six measurement time points; however, this analysis focuses solely on the first two laboratory sessions (T0 and T1), during which a fear conditioning experiment was conducted. Participants received a financial compensation of 170 Euro for completing the sessions at T0 and T1.

From the original sample, 97 participants ($female_{T1}$ = 64, $male_{T1}$ = 64 and 33, age $_{M,T1}$ = 25.20, $age_{SD,T1}$ = 3.73, $age_{range,T1}$ = 19 - 35) returned to the follow up measurement after six months (T1). Prior to data analysis, several participants had to be excluded (see Supplementary Figure 1 for exclusion details). Finally, data of 69 participants were analyzed.

Supplementary Table 1: Descriptive information on the subsamples being exposed or unexposed to recent adversity.

| Variable | Recent adversity prior to T1 | No recent adversity prior to T1 | Statistics |
|---|---|---|---|
| **N** | 24 (35%) | 45 (65%) | $X^2$(1) = 6.39, p = .011 |
| **Female/Male** | 17 (71%) / 7 (29%) | 23 (51%) / 22 (49%) | $X^2$ (1) = 1.75, p = .185 |
| **Recent adversity at T0 (yes/no)** | 14 (58%) / 10 (42%) | 14 (31%) / 31 (69%) | $X^2$ (1) = 3.75, p = .053 |
| **Unaware or semi-aware/aware at T0** | 6 (25%) / 18 (75%) | 7 (16%) / 38 (84%) | $X^2$ (1) = 0.40, p = .527 |
| **Unaware or semi-aware/aware at T1** | 2 (8%) / 22 (92%) | 6 (13%) / 39 (87%) | $X^2$ (1) = 0.05, p = .823 |
| **Age (M/SD)** | 25.12 ( 3.85) | 24.71 ( 3.80) | t(67) = -0.43, p = 0.669, d = -0.11 |
| **STAI-T sum (M/SD)** | 37.21 ( 7.77) | 33.82 ( 7.06) | t(43.3) = -1.78, p = 0.082, d = -0.46 |
| **STAI-S sum at T0 day1 (M/SD)** | 35.50 ( 5.38) | 34.86 ( 4.30) | t(39.8) = -0.5, p = 0.619, d = -0.14 |
| **STAI-S sum at T0 day2 (M/SD)** | 36.96 ( 7.29) | 34.84 ( 6.53) | t(67) = -1.23, p = 0.223, d = -0.31 |
| **STAI-S sum at T1 day1 (M/SD)** | 38.30 ( 6.10) | 35.30 ( 5.81) | t(65) = -1.98, p = 0.052, d = -0.51 |

| Variable | Recent adversity prior to T1 | No recent adversity prior to T1 | Statistics |
|---|---|---|---|
| **STAI-S sum at T1 day2 (M/SD)** | 38.50 ( 7.71) | 34.31 ( 6.30) | t(67) = -2.43, p = 0.018, d = -0.61 |
| **US intensity at T0 (M/SD in mA)** | 11.03 (12.53) | 8.25 ( 7.61) | t(67) = -1.15, p = 0.255, d = -0.29 |
| **US intensity at T1 (M/SD in mA)** | 11.73 (17.86) | 8.98 (10.25) | t(67) = -0.82, p = 0.418, d = -0.21 |
| **US aversiveness at T0 day 1 (M/SD, VAS: 0-25)** | 21.09 ( 2.27) | 19.52 ( 3.20) | t(60) = -2.03, p = 0.047, d = -0.54 |
| **US aversiveness at T0 day 2 (M/SD, VAS: 0-25)** | 15.52 ( 4.71) | 17.14 ( 3.31) | t(64) = 1.63, p = 0.109, d = 0.42 |
| **US aversiveness at T1 day 1 (M/SD, VAS: 0-25)** | 19.62 ( 3.03) | 20.29 ( 2.89) | t(64) = 0.88, p = 0.384, d = 0.22 |
| **US aversiveness at T1 day 2 (M/SD, VAS: 0-25)** | 16.58 ( 4.22) | 18.55 ( 3.57) | t(64) = 1.63, p = 0.109, d = 0.42 |

Note. Awareness relates to the awareness of CS-US contingencies. Semi-aware means that it was unclear if participants were aware or not. STAI-T data were derived from the study participants were preselectet from. M = mean, SD = standard deviation, STAI-T = State-Trait Anxiety Inventory, Trait scale (4), STAI-S = State-Trait Anxiety Inventory, State scale (4). Due to alternative recruitment procedures, STAI-T was assessed 1–9 months prior to T0. US = unconditioned stimulus.

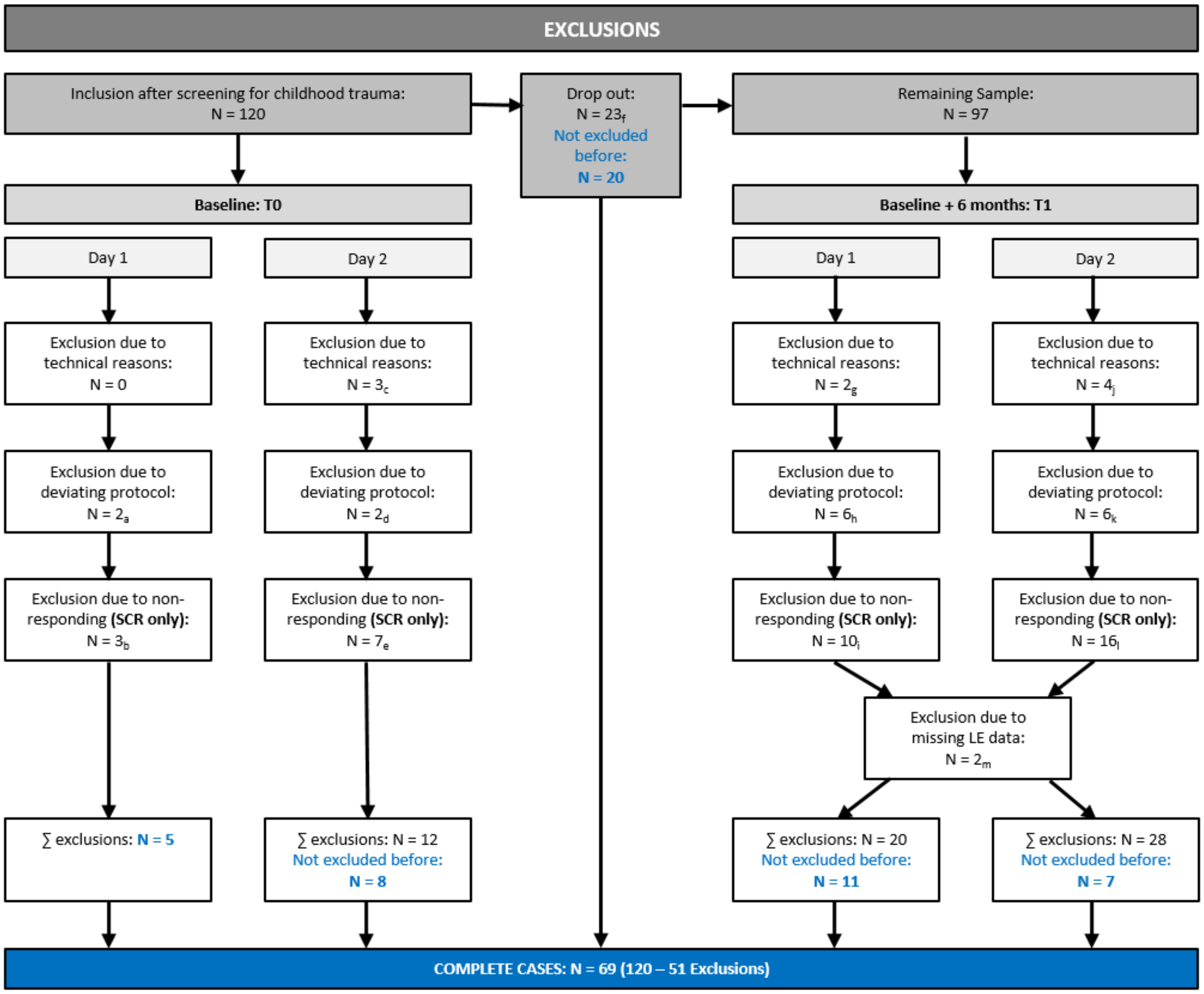


Supplementary Figure 1: Overview of sample exclusions.

#### Stimuli

Visual stimuli were identical for all participants, but allocation to CS+/CS- and CS-type of the first trial of each phase was counterbalanced between participants. Conditioned stimuli were two light grey fractals (RGB [230,230,230]), 492*492 pixel) and presented in a pseudo-randomized order for 6 - 8 s (mean: 7 s) with a maximum of two consecutive identical stimuli. A white fixation cross served as ITI and was shown for 10 - 16 s (mean: 13 s). All stimuli were presented by the software Presentation (Version 14.8, Neurobehavioral Systems, Inc., Albany California, USA). All stimuli were presented on a grey background (RGB [100, 100, 100]) keeping the context constant aiming to avoid renewal effects during the reinstatement manipulation (5).

The electrotactile stimulus, serving as US, consisted of three 2 ms rectangular pulses (interpulse interval: 50 ms, onset: 0.2 s before CS+ offset) and was administered to the back of the right hand of participants. It was generated by a Digitimer DS7A constant current stimulator (Welwyn Garden City, Hertfordshire, UK) and delivered through a 1 cm diameter platinum pin surface electrode (Speciality Developments, Bexley, UK). The electrode was attached between the metacarpal bones of the index and middle finger. US intensity was individually calibrated in a standardized step-wise procedure controlled by the experimenter. The aim was an unpleasant, but still tolerable level rated by participants between seven and eight on a scale from zero (= stimulus was not unpleasant at all) to ten (= stimulus was the worst one could imagine within the study context). Participants were, however, not informed that we aimed at a score of seven to eight.

**Experimental Design and Procedure**

This study was part of a larger longitudinal study that spanned six time points. Data from a differential 2-day fear conditioning experiment, collected at two time points (T0 and T1 6 months later) as well as questionnaire data from three online follow-ups after 1, 1.5 and 2 years following T1 (T2-T4) are reported here. At both time points, participants underwent an identical paradigm in the MR environment with habituation and fear acquisition training on day 1 and extinction training, reinstatement and reinstatement-test on day 2.

After filling in the informed consent on day 1, participants completed a paper version of the STAI-S and a rating training. Participants were positioned inside the scanner and equipped with SCR electrodes, the electrode for electrotactile stimulation and a response button box. To ensure wakefulness of participants, eye tracking was calibrated using EyeLink 1000 Desktop Mount systems (SR Research Ltd., Mississauga, Ontario, Canada). Both eyes were recorded at 1000 Hz. Subsequent to structural scans and the US calibration (see section Stimuli), habituation and acquisition training took place. Habituation and acquisition training consisted of 7 and 14 presentations of each CS (i.e. CS+ and CS-) respectively. During habituation, CSs were presented without US occurrence, which was communicated to the participants. During the uninstructed acquisition training phase, 100% of the CS+ presentations were reinforced (i.e., participants were not explicitly informed about CS-US contingency). Assessments on day 1 ended with the postexperimental awareness interview (see section ‘CS-US contingency awareness’). Overall, three saliva samples were taken on day 1 with one sample before attachment of the electrodes (sample 1) and two samples after acquisition training (sample 2: 20 min after sample 1; sample 3: 30 min after sample 1).

Rating training and US re-calibration were not repeated on day 2. The procedure on day 2 included extinction training, reinstatement and reinstatement-test. Extinction training consisted of 28 trials for each CS (14 CS+/14 CS−), and the reinstatement-test phase of 14 trials each CS (7 CS+/7 CS−). Reinstatement consisted of three trials with a duration of 5 s each presented after a 10 s ITI. Reinstatement USs were delivered 4.8 s after each trial onset. The reinstatement phase was followed by a 13 s ITI before the next CS was presented during reinstatement-test. After the postexperimental interview, participants filled in a battery of questionnaires including questions on life events (see sections ‘Additional questionnaires’ and ‘Quantification of life adversity’, respectively) computerized using LimeSurvey software (LimeSurvey GmbH, Hamburg,

Germany, URL http://www.limesurvey.org). Overall, five saliva samples were extracted on day 2: before attachment of the electrodes (sample 1), after RI-test (sample 2: 30 min after sample 1), subsequent to structural scans (sample 3: 50 min after sample 1), and as well as 60 min and 70 min after sample 1 (i.e., sample 4 and 5).

For hair cortisol analyses, hair samples (at least 3cm) were collected on day 1 or day 2 at T0 and T1 and also at an additional visit approximately 3 months after the first extraction (T0) to cover the entire 6 month interval.

**Skin Conductance Responses**

SCRs were recorded continuously with a 1000 Hz sampling rate using a BIOPAC MP 100 amplifier and AcqKnowledge 3.9.2 software (BIOPAC Systems, Inc., Goleta, California, USA). For analog to digital conversion, a CED2502-SA with Spike 2 software (Cambridge Electronic Design, Cambridge, UK) was used. Two self-adhesive SCR electrodes (Ø = 55 mm) with hydrogel and Ag/AgCl-sensor recording were attached on the palm of the left hand, i.e., on the distal and proximal hypothenar. A 1 Hz lowpass filter and a gain of 5 or 10 μΩ were applied.

SCRs were scored semi-manually (i.e., computer assisted) by using a custom-made computer program (EDA View, provided by Prof. Dr. Matthias Gamer, University of Würzburg) as the first response from trough to peak 0.9 – 3.5 s after CS onset (0.9 – 2.5 s after US onset, 6). The maximum rise time was set to 5 s (7). Data were downsampled to 10 Hz. Amplitudes below 0.01 μS were classified as zero-responses (8). Data with recording artifacts or excessive baseline activity were treated as missing data points and excluded from the analyses. The response (i.e., trough-to-peak) suggested by the scoring algorithm were inspected visually, and adjusted if necessary. Scoring was performed blind to stimulus type (i.e., CS+, CS-, US). Raw SCR amplitudes were log-transformed by taking the natural logarithm and range corrected by dividing each log-transformed SCR through the maximum amplitude per participant and day (CSs or US) (9,10). Due to potentially different response ranges, the range correction was applied for the two days separately (8).

On day 1, participants with more than 2/3 zero-responses to the US during fear acquisition were classified as physiological non-responders and excluded from analyses (8). On Day 2, participants were classified as non-responders if they showed no response to any of the three reinstatement USs (see Supplementary Figure 1).

**Fear Ratings**

Fear ratings were provided after habituation and acquisition, before and after extinction as well as after reinstatement-test. Participants were asked on the stimulus screen how much "stress, fear, and tension" they experienced, when they saw the CS+ and CS– the last time. After reinstatement-test, both CSs were additionally rated retrospectively referring to their first presentation following the reinstatement. Answers were given within 5 s on a visual analog scale (VAS), which ranged from zero (answer = none) to 100 (answer = maximum) by using the button box. Pressing the buttons moved a bar on the VAS to the aimed value and also logged in the answers. Ratings had to be confirmed via button press. Not registered ratings were considered as missing values. On both days, the same procedure was used to rate the unpleasantness of the electrotactile stimulus. For analyses, the rating scale was reduced to 0 - 25.

**CS-US Contingency Awareness**

After acquisition training on day 1, a standardized post-experimental awareness interview (adapted from 11) was conducted. Based on the provided answers, the experimenter classified the participants as aware, unaware, or uncertain of CS-US contingencies.

**Quantification of Recent Adversity**

Recent life adversity was assessed by a modified version of the Life Events Checklist (LEC, 12,13) as in (14), whose findings we aimed to replicate here in a longitudinal setting. However, inspection of the raw data revealed invalid completion. Participants completed the questionnaire online and answers to questions about the valence (positive, negative, irrelevant) and date of events were in an open format. Unfortunately, we could not utilize the LEC because participants' responses were inconclusive, raising concerns about data quality. Therefore, an adapted version of the List of Threatening Experiences (LTE, 15), which was also recorded, was used to quantify recent life adversity. The questionnaire captures the exposition to twelve different life events during the last six months comprising illness or injury to self or close relatives, death of relatives or friends, separation after marriage, broke of a steady relationship, serious problems with close persons, unemployment for > 1 month, sacked from job, serious financial crisis, problems with police or court appearance and loss or theft of something valuable. Test-retest reliability and concurrent validity of the original English version were reported as good (16).

**Additional Questionnaires**

After the post-experimental interview on day 2, the participants completed a battery of questionnaires, always presented in the identical order, computerized via LimeSurvey: a short questionnaire for the assessments of stressors (Kurzer Fragebogen zur Erfassung von Belastungen, KFB, 17), the german versions of the social support appraisals scale (SS-A-d, 18), the Trier Inventory for Chronic Stress (TICS, 19), the stress coping questionnaire (Stressverarbeitungsfragebogen, SVF 78, 20), Berliner Social-Support Scales (BSSS, 21), Coping Orientation to Problems Experienced Inventory (Brief COPE, 22), Cognitive Emotion Regulation Questionnaire (CERQ short, 23), Generalized Self-Efficacy Scale (GSE, 24), Life Events Checklist (LEC, modified version, 12,13), Perceived Stress Questionnaire (PSQ, 25) and self-constructed questions relevant for cortisol. STAI-T scores were acquired 1–9 months prior to this study and are reported here for completeness. Given its established temporal stability for short time intervals, we regard the STAI-T as an appropriate trait indicator (26). During the follow-up period, participants completed online questionnaires at three follow-up time points (T2–T4), corresponding to 1-, 1.5-, and 2-year follow-ups after T1, using the State-Trait Anxiety Inventory (STAI-T, 4) and Beck Depression Inventory-II (BDI-II, 27) to assess anxiety and depression levels.

**Salivary Cortisol**

Saliva was extracted by asking the subjects to move the suction roll of Salivette devices (SARSTEDT AG & Co., Nümbrecht, Germany) in the mouth for 60 s and put it into the appendant tube. Saliva samples were kept at room temperature until the end of the session and shelved afterwards at -20 degrees C. They were shipped to and analysed by the Dresden LabService GmbH, Dresden, Germany (http://www.labservice-dresden.de). Prior to analysis, Salivettes were defreezed and centrifuged for 5 min at 3000 rpm. This resulted in a clear supernatant featuring low viscosity. For measuring salivary concentrations, chemiluminescence immunoassay with high sensitivity (IBL International, Hamburg, Germany) was used. Intra and interassay cortisol coefficients fell below 8%. Since salivary cortisol data were positively skewed, individual values were log-transformed. Visual inspection indicated that the transformation successfully normalized the distribution.

**Hair Cortisol**

Hair strands were cut carefully from a posterior vertex position near the scalp by using fine scissors. The specific number of extracted strands varied, depending on the participants' consent to cut more or less hair. For analysis in the laboratory (Dresden LabService GmbH, Dresden, Germany, http://www.labservice-dresden.de), hair strands were lined up and segmented in slices of 3 cm. The protocol of (28) was applied for the following hair washing and steroid extraction. Briefly, after putting each hair segment together with 2.5 ml isopropanol in a 10 ml glass container, it was mixed for 3 min in an overhead rotator. The glass container was decanted and the washing procedure was repeated two times. After drying hair segments for 12 hours, they were weighed and 7.5 mg were put into a 2 ml cryovial together with 1.5 ml pure methanol. In the next step, steroid extraction was conducted for 18 hours. Hair samples were centrifuged in a microcentrifuge for 2 min at 10000 rpm and 1 ml of the resulting clear supernatant was put into a fresh 2 ml glass vial. Afterwards, they were entirely dried by evaporating the alcohol at 50 degrees C under a constant nitrogen stream. The vial was vortexed for 15 s after adding 0.4 ml of water. To determine cortisol, 50 ml were taken from the tube and an immunoassay with chemiluminescence detection (CLIA, IBL-Hamburg, Germany) was applied. Intra and interassay cortisol coefficients of assay variance fell below 8%. Hair cortisol concentrations are often log-transformed to address non-normality. However, inspection of model assumptions - including residual normality, homoscedasticity, and the distribution of random intercepts - indicated that the untransformed data provided a better fit to these assumptions than the log-transformed values.

**Statistical Analyses**

For analyses of SCRs during fear acquisition training, the dependent variable consisted of the mean of trials per CS type except the first trial. First CS trials were excluded, because conditioning could not have taken place yet because the SCR to the first CS+ is recorded prior to the first US presentation. The first trial of the 24h-delayed extinction training phase was analyzed separately as “fear recall”. For the analysis of fear recall, the last trial of acquisition training and the first trial of extinction training were included, while for reinstatement-test, the last trial of extinction training and the first trial of reinstatement-test were included (5). Because SCRs to the US were aggregated at the participant level, eliminating within-subject dependence, a linear model was used instead of linear mixed-effects model with SCRs to the US at T1 as the dependent variable, recent adversity as the independent variable, and SCRs to the US at T0 as a covariate.

For analysis of fear ratings during acquisition training, the dependent variable consisted of the pre- and post rating for acquisition and the pre- as well as post-extinction ratings for extinction training, respectively. To analyze fear recall, the post-acquisition and the pre-extinction ratings were entered into the analysis, while for analyzing reinstatement, the post-extinction rating as well as the pre-reinstatement-test rating were included.

Prediction of anxiety and depression levels was examined using linear regression analyses, with differential SCRs and extracted neural parameter estimates from the CS+ versus CS− contrast during acquisition, fear recall, and reinstatement serving as predictors, and STAI-T and BDI-II scores at T2-T4 as outcome variables in separate models. We restricted the analyses to phases and outcome measures for which significant effects were observed. Accordingly, linear regression analyses were not conducted for extinction training or for rating measures.

Hair cortisol levels across sampling points were analyzed using linear mixed-effects models. Trial and life-event exposure were entered as fixed effects, along with their interaction. Baseline hair cortisol levels (T0) were included as a covariate and were centered within individuals. Random intercepts were specified for subjects.

For linear mixed-effects models, significance of fixed effects was evaluated using F-tests derived from model-based ANOVA tables, with denominator degrees of freedom approximated by Satterthwaite's method. Post hoc pairwise comparisons of estimated marginal means were conducted, with Holm correction applied to adjust for multiple testing.

Sample characteristics, including age and sex, as well as experiment-related measures, including awareness of CS–US contingencies and US-evoked SCRs, were compared between individuals exposed and unexposed to RA using two-sample t-tests and chi-square tests, as appropriate (see Supplementary Table 1).

For data analysis and visualizations as well as for the creation of the manuscript, we used R (Version 4.4.1; 29) and the R-packages *afex* (30), *apa* (Version 0.3.4; 31,32), *car* (Version 3.1.2; 33), *carData* (Version 3.0.5; 34), *cowplot* (Version 1.1.3; 35), *dplyr* (Version 1.1.4; 36), *effects* (Version 4.2.2; 37-39), *effectsize* (Version 0.8.9; 40), *effsize* (Version 0.8.1; 41), *emmeans* (Version 1.10.4; 42), *ez* (Version 4.4.0; 43), *flextable* (Version 0.9.6; 44), *forcats* (Version 1.0.0; 45), *ggExtra* (Version 0.10.1; 46), *ggplot2* (Version 4.0.3; 47), *ggpubr* (Version 0.6.0; 48), *knitr* (Version 1.48; 49), *lme4* (Version 1.1.35.5; 50), *lmerTest* (Version 3.1.3; 51), *lubridate* (Version 1.9.3; 52), *Matrix* (Version 1.7.0; 53), *MESS* (Version 0.5.12; 54), *nlme* (Version 3.1.168; 55), *papaja* (Version 0.1.2; 56), *patchwork* (Version 1.3.2; 57), *performance* (Version 0.12.3; 58), *psychReport* (Version 3.0.2; 59), *purrr* (Version 1.0.2; 60), *readr* (Version 2.1.5; 61), *readxl* (Version 1.4.3; 62), *rstatix* (Version 0.7.2; 63), *sjPlot* (Version 2.8.16; 64), *stringr* (Version

1.5.1; 65), *tibble* (Version 3.2.1; 66), *tidyr* (Version 1.3.1; 67), *tidyverse* (Version 2.0.0; 68), and *tinylabels* (Version 0.2.4; 69).

**fMRI Data Acquisition, Preprocessing, and Statistical Analysis**

MRI data were acquired on a 3 Tesla MR-scanner (PrismaFit, Siemens, Germany) using a 64-channel head coil. Functional data were obtained using an echo planar images sequence (TR=1980 ms, TE=30 ms). For each volume, 54 slices with a voxel size of 1.7 x 1.7 x 1.7mm (0.5 mm gap) were acquired sequentially. Structural images were obtained by using a T1-weighted MPRAGE sequence. fMRI data were analyzed using SPM12 (Welcome Trust Centre for Neuroimaging, UCL, London, UK). Preprocessing included coregistration to the individual structural image, realignment, normalization to group-specific templates created via the DARTEL-algorithm (70) as well as smoothing (6mm FWHM). For brain images, we used Synth-Strip for skull stripping (71).

At the first level for acquisition and extinction training, four effects-of-interest regressors (i.e., CS+ and CS- at T0 and T1, respectively) averaged across trials as well as several nuisance (i.e., start screen of experiment, ratings, four linear time modulators for CS+ and CS-, six movement parameters derived from realignment - each for both time points) were built. For acquisition training, US and omitted US were also included as regressors. For the first levels for fear recall and reinstatement-test, we followed a single trial approach across all phases integrating all CS+ and CS- trials as effects-of-interest regressors. In addition, we integrated all nuisance regressors as listed for the acquisition and extinction training first levels plus the US during reinstatement as well as omitted US. Single trials were used due to the temporal transience of the reinstatement-effect (5).

All regressors of interest were modeled as stick function and time locked to stimulus onset (CS/US). Regression coefficients (beta values) for the regressor in each voxel were computed via the general linear model. Contrasts of interest (CS+ > CS−) were estimated on the first level and taken to the second-level analyses where interaction effects between RA group and Time point were assessed using a flexible factorial design implemented in SPM. T-contrasts were specified with contrast weights [1 −1 −1 1] and [−1 1 1 −1]. The first contrast, (i.e., [1 −1 −1 1]) tested for lower CS discrimination (i.e., CS+ > CS-) in the unexposed vs. exposed group at T1 relative to T0, while the second contrast (i.e., [−1 1 1 −1]) tested for higher CS discrimination in the unexposed vs. exposed group at T1 relative to T0. Moreover, predictions of anxiety and depression levels were tested by using linear regression models with extracted neural parameter estimates from the CS+ versus CS− contrast for acquisition, fear recall, and reinstatement as predictors and STAI-T/BDI-II scores at T2 - T4 as outcomes.

Small volume corrections were performed for predefined regions of interest (ROIs), based on key areas implicated in previous studies (72,73). Anatomical ROIs for the bilateral amygdala, hippocampus, caudate nucleus, putamen, pallidum, nucleus accumbens, and thalamus were derived from the Harvard–Oxford atlas (74) using a maximum probability threshold of 0.5. The anterior insula ROI was defined as the intersection between the 0.5-thresholded Harvard–Oxford mask and a 60 × 30 × 60 mm box centered at MNI coordinates (0, 30, 0), following anatomical subdivisions described in (75). Coordinate-based ROIs for the dlPFC and dACC were defined as 20 × 16 × 16 mm boxes centered on peak coordinates from a meta-analysis (76). The vmPFC ROI was defined using coordinates from previous fear-conditioning studies (77,78). Coordinates were: left dlPFC (−36, 44, 22), right dlPFC (34, 44, 32), dACC (0, 18, 42), and vmPFC (0, 40, −12). For midline regions (dACC and vmPFC), the x-coordinate was set to 0 to ensure symmetry. For all fMRI analyses a peak voxel, familywise error (FWE) corrected, threshold at $p < 0.05$ was considered significant. Unthresholded T-maps for CS+ > CS- contrasts are available on NeuroVault (https://identifiers.org/neurovault.collection:24143).

#### Results

##### Salivary cortisol

Absolute cortisol levels across sampling points were analyzed using linear mixed-effects models, fitted separately for each experimental day. Time point within experimental day and RA exposure were specified as fixed effects, including their interaction. Baseline mean cortisol levels were included as a covariate and were centered within individuals. Random intercepts were specified for subjects to account for repeated measurements.

For salivary cortisol levels on day 1 and day 2, no significant main effect of or interaction with RA were observed. However, there was a significant main effect of Time at both days indicating significantly decreasing salivary cortisol levels across time points (see Figure see Supplementary Figure 2; day 1: $F(2{,}117.33) = 44.48$, $p < .001$, day 2: $F(4{,}240.28) = 29.68$, $p < .001$), with concentrations at all time points within the experimental day differing from each other on day 1 (Time 1 vs. 2: $t(117.30) = 5.58$, $p_{\mathrm{Holm}(3)} < .001$, Time 1 vs. 3: $t(117.59) = 9.41$, $p_{\mathrm{Holm}(3)} < .001$, Time 2 vs. 3: $t(117.27) = 5.57$, $p_{\mathrm{Holm}(3)} < .001$), but with a few exceptions on day 2 (Time 2 vs. 3: $t(240.12) = 2.15$, $p_{\mathrm{Holm}(10)} = .066$, Time 4 vs. 5: $t(240.36) = 1.69$, $p_{\mathrm{Holm}(10)} = .091$). On day 2, saliva cortisol concentration at T1 was also significantly related to that of T0, $F(1{,}240.26) = 14.63$, $p < .001$.

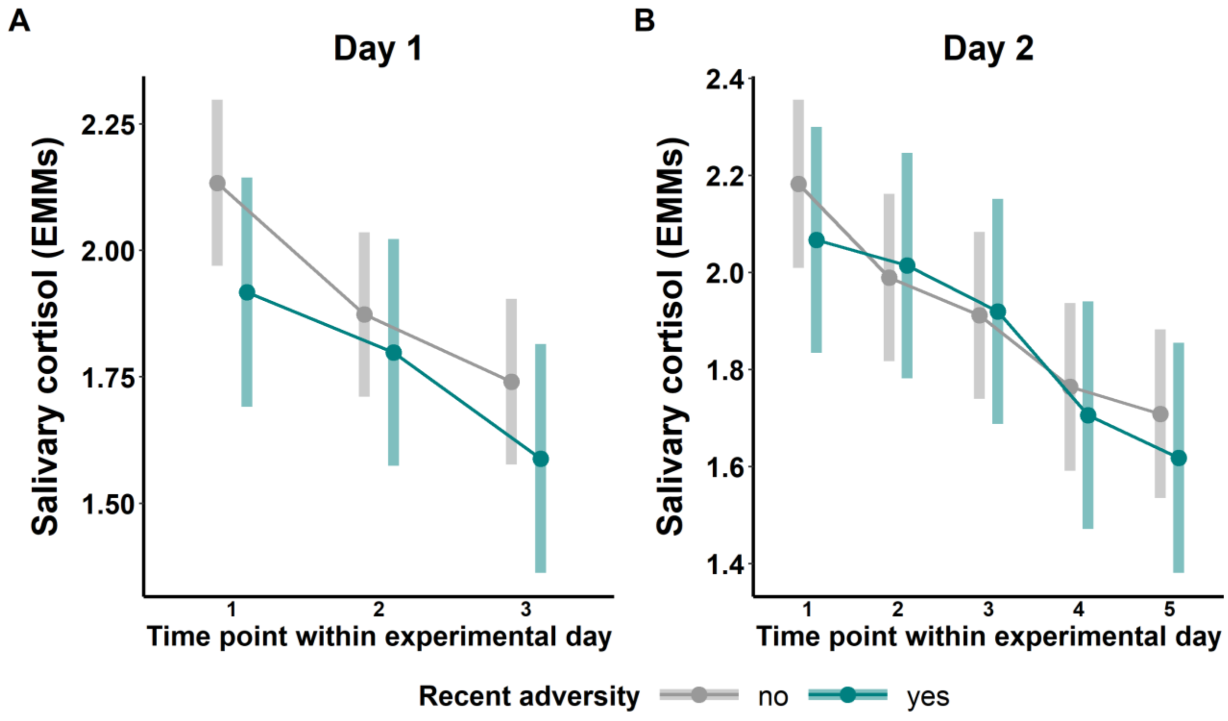


Supplementary Figure 2. Estimated marginal means (EMMs) of salivary cortisol concentrations across different time points (upper row) on day 1 (A) and day 2 (B).

**Hair cortisol**

For hair cortisol, there were no significant main effects or interactions involving Group (see Supplementary Table 2). However, we observed a significant main effect Time, $F(5{,}238.37) = 4.14$, $p = .001$. Post hoc comparisons showed that hair cortisol concentration was significantly higher during month 2 vs. 5 ($t(242.40) = 3.21$, $p_{\text{Holm}(15)} = .020$), during month 2 vs. 6 ($t(242.11) = 3.41$, $p_{\text{Holm}(15)} = .011$) as well as during month 3 vs. 5 ($t(241.95) = 3.13$, $p_{\text{Holm}(15)} = .023$) and during 3 and 6 ($t(241.63) = 3.35$, $p_{\text{Holm}(15)} = .013$). Furthermore, there was a significant main effect of hair cortisol level at T0, suggesting a positive association between baseline (T0) levels and hair cortisol at later time points, $F(1{,}66.89) = 11.87$, $p < .001$.

Supplementary Table 2. Complete results of the mixed-effects model analyzing hair cortisol levels.

| | Hair_cortisol_level | | | |
|---|---|---|---|---|
| *Predictors* | *Estimates* | *CI* | *p* | *df* |
| (Intercept) | 1.64 | 1.20 – 2.07 | **<0.001** | 63.55 |
| Month after T0 [1] | -0.00 | -0.15 – 0.14 | 0.953 | 233.21 |
| Month after T0 [2] | 0.19 | 0.05 – 0.34 | **0.010** | 233.81 |
| Month after T0 [3] | 0.16 | 0.01 – 0.31 | **0.036** | 234.28 |
| Month after T0 [4] | -0.09 | -0.22 – 0.05 | 0.217 | 233.15 |
| Month after T0 [5] | -0.15 | -0.29 – -0.01 | **0.035** | 233.49 |
| Recent adversity [1] | 0.13 | -0.16 – 0.41 | 0.374 | 68.28 |
| Hair cortisol level T0 | 0.30 | 0.13 – 0.47 | **0.001** | 66.75 |
| Month after T0 [1] × Recent adversity [1] | 0.05 | -0.30 – 0.40 | 0.773 | 238.70 |
| Month after T0 [2] × Recent adversity [1] | 0.03 | -0.29 – 0.34 | 0.865 | 237.37 |
| Month after T0 [3] × Recent adversity [1] | 0.06 | -0.25 – 0.36 | 0.720 | 237.42 |
| Month after T0 [4] × Recent adversity [1] | 0.02 | -0.23 – 0.27 | 0.875 | 238.32 |
| Month after T0 [5] × Recent adversity [1] | -0.01 | -0.26 – 0.23 | 0.918 | 242.02 |
| **Random Effects** | | | | |
| $\sigma^2$ | 0.22 | | | |
| $\tau_{00\ Subject}$ | 0.22 | | | |
| ICC | 0.50 | | | |
| $N_{Subject}$ | 63 | | | |
| Observations | 304 | | | |
| Marginal $R^2$ / Conditional $R^2$ | 0.119 / 0.558 | | | |

#### Manipulation check

Supplementary Table 3: Manipulation check for SCR and fear ratings.

| Measure | Phase | Timepoint | *t* | *df* | *p* | *Cohen's d* | *LL (95% CI)* | *UL (95% CI)* |
|---|---|---|---|---|---|---|---|---|
| SCR | Acquisition Training | T0 | **11.19** | **68** | **0.000** | **0.95** | **0.75** | **1.15** |
| | | T1 | **4.04** | **109** | **0.000** | **0.96** | **0.61** | **1.31** |
| | Extinction Training | T0 | 1.06 | 136 | 0.292 | 0.18 | -0.16 | 0.51 |
| | | T1 | 1.29 | 136 | 0.198 | 0.25 | -0.09 | 0.58 |
| Fear Ratings | Acquisition Training | T0 | **14.18** | **116** | **0.000** | **3.07** | **2.57** | **3.57** |
| | | T1 | **23.37** | **104** | **0.000** | **5.89** | **5.11** | **6.66** |
| | Extinction Training | T0 | **2.98** | **119** | **0.003** | **0.63** | **0.28** | **0.97** |
| | | T1 | **4.11** | **105** | **0.000** | **1.01** | **0.65** | **1.36** |

Note. Bold numbers indicate significant results ($p < 0.05$).

#### Comprehensive SCR Results

Supplementary Table 4. Complete results of the mixed-effects models analyzing SCRs.

| | SCR_ACQ | | | | SCR_EXT | | | | SCR_FREC | | | | SCR_RI | | | |
|---|---|---|---|---|---|---|---|---|---|---|---|---|---|---|---|---|
| *Predictors* | *Estimates* | *CI* | *p* | *df* | *Estimates* | *CI* | *p* | *df* | *Estimates* | *CI* | *p* | *df* | *Estimates* | *CI* | *p* | *df* |
| (Intercept) | 0.17 | 0.13 – 0.22 | **<0.001** | 67.00 | 0.06 | 0.04 – 0.08 | **<0.001** | 67.00 | 0.19 | 0.14 – 0.23 | **<0.001** | 67.29 | 0.18 | 0.14 – 0.22 | **<0.001** | 67.00 |
| CS type1 | 0.05 | 0.02 – 0.07 | **0.001** | 66.00 | 0.01 | 0.00 – 0.02 | **0.008** | 66.00 | 0.07 | 0.04 – 0.09 | **<0.001** | 196.69 | 0.05 | 0.02 – 0.09 | **0.001** | 200.00 |
| Recent adversity [1] | -0.08 | -0.15 – -0.01 | **0.027** | 67.00 | -0.03 | -0.07 – 0.01 | 0.112 | 67.00 | -0.05 | -0.13 – 0.03 | 0.204 | 66.56 | -0.04 | -0.11 – 0.02 | 0.192 | 67.00 |
| SCR T0 | 0.22 | -0.03 – 0.46 | 0.079 | 66.00 | 0.28 | 0.10 – 0.46 | **0.003** | 66.00 | 0.03 | -0.09 – 0.15 | 0.590 | 198.25 | 0.20 | 0.04 – 0.36 | **0.012** | 200.00 |
| CS type1 × Recent adversity [1] | -0.03 | -0.06 – -0.00 | **0.033** | 66.00 | -0.01 | -0.02 – 0.00 | 0.228 | 66.00 | -0.06 | -0.10 – -0.01 | **0.016** | 196.29 | -0.05 | -0.11 – 0.00 | 0.052 | 200.00 |
| Time1 | | | | | | | | | -0.07 | -0.10 – -0.04 | **<0.001** | 196.97 | -0.08 | -0.12 – -0.04 | **<0.001** | 200.00 |
| Time1 × CS type1 | | | | | | | | | -0.02 | -0.04 – 0.01 | 0.217 | 196.66 | -0.02 | -0.05 – 0.01 | 0.265 | 200.00 |
| Time1 × Recent | | | | | | | | | 0.01 | -0.04 – 0.05 | 0.703 | 196.95 | 0.01 | -0.04 – 0.06 | 0.701 | 200.00 |

| | | | | | | | | | | | | | | | | |
|---|---|---|---|---|---|---|---|---|---|---|---|---|---|---|---|---|
| adversity [1] | | | | | | | | | | | | | | | | |
| (Time1 × CS type1) × Recent adversity [1] | | | | | | | | | 0.01 | -0.03 – 0.06 | 0.617 | 196.28 | 0.01 | -0.04 – 0.07 | 0.608 | 200.00 |
| **Random Effects** | | | | | | | | | | | | | | | | |
| $\sigma^2$ | 0.01 | | | | 0.00 | | | | 0.03 | | | | 0.05 | | | |
| $\tau_{00}$ | 0.02 Subject | | | | 0.01 Subject | | | | 0.02 Subject | | | | 0.01 Subject | | | |
| ICC | 0.72 | | | | 0.85 | | | | 0.33 | | | | 0.11 | | | |
| N | 69 Subject | | | | 69 Subject | | | | 69 Subject | | | | 69 Subject | | | |
| Observations | 138 | | | | 138 | | | | 272 | | | | 276 | | | |
| Marginal $R^2$ / Conditional $R^2$ | 0.175 / 0.770 | | | | 0.057 / 0.862 | | | | 0.148 / 0.433 | | | | 0.239 / 0.325 | | | |

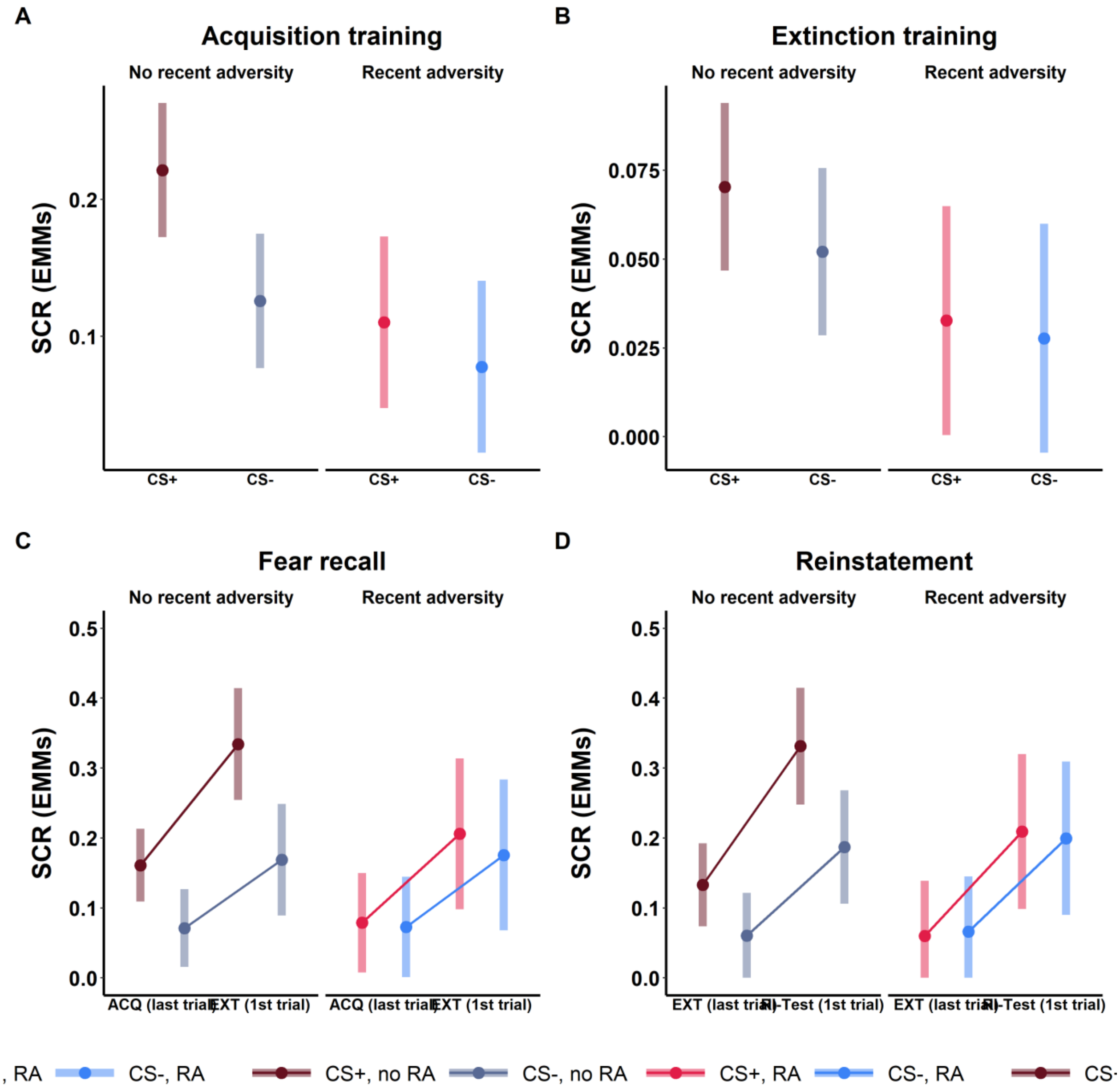


Supplementary Figure 3: Estimated marginal means (EMMs) of SCRs at T1 (controlled for T0 responding) during acquisition training (A), extinction training (B), for fear recall (C) and reinstatement (D) separated by CS type and exposure to recent adversity. CS = conditioned stimulus. ACQ = acquisition training, EXT = extinction training, RI = reinstatement.

**Prediction of anxiety and depression levels**

Supplementary Table 5: Prediction of anxiety and depression levels by CS discrimintion in SCR.

| Phase | Criterion | Timepoint | *beta* | *SE beta* | *t* | *p* | *R squared* | *Cohen's f2* |
|---|---|---|---|---|---|---|---|---|
| **Acquisiton Training** | **STAI-T** | **T3** | -1.17 | 11.31 | -0.10 | 0.918 | 0.00 | 0 |
| | | **T4** | -0.88 | 11.73 | -0.08 | 0.940 | 0.00 | 0 |
| | | **T5** | 3.30 | 11.10 | 0.30 | 0.767 | 0.00 | 0 |
| | **BDI** | **T3** | -8.05 | 7.20 | -1.12 | 0.267 | 0.02 | 0 |
| | | **T4** | **-17.60** | **8.36** | **-2.10** | **0.039** | **0.07** | **0** |
| | | **T5** | -4.71 | 6.57 | -0.72 | 0.476 | 0.01 | 0 |
| **Fear Recall** | **STAI-T** | **T3** | -2.22 | 4.74 | -0.47 | 0.641 | 0.00 | 0 |
| | | **T4** | -0.80 | 5.16 | -0.15 | 0.878 | 0.00 | 0 |
| | | **T5** | 5.96 | 4.73 | 1.26 | 0.213 | 0.03 | 0 |
| | **BDI** | **T3** | **-6.24** | **3.02** | **-2.07** | **0.043** | **0.06** | **0** |
| | | **T4** | 1.10 | 3.80 | 0.29 | 0.773 | 0.00 | 0 |
| | | **T5** | 5.17 | 2.81 | 1.84 | 0.071 | 0.05 | 0 |
| **Reinstatement** | **STAI-T** | **T3** | -3.54 | 3.62 | -0.98 | 0.332 | 0.01 | 0 |
| | | **T4** | -3.47 | 3.71 | -0.94 | 0.353 | 0.01 | 0 |
| | | **T5** | -1.09 | 3.48 | -0.31 | 0.756 | 0.00 | 0 |
| | **BDI** | **T3** | 0.53 | 2.34 | 0.23 | 0.822 | 0.00 | 0 |
| | | **T4** | 0.01 | 2.76 | 0.00 | 0.997 | 0.00 | 0 |
| | | **T5** | 1.30 | 2.06 | 0.63 | 0.531 | 0.01 | 0 |

Note. STAI-T = State-Trait Anxiety Inventory, Spielberger et al., 1983; BDI = Beck Depression Inventory-II, Beck et al., 1996. Bold numbers indicate significant results ($p < 0.05$).

Supplementary Table 6: Prediction of anxiety and depression levels by CS discrimintion in BOLD fMRI.

| **Phase** | **Region** | **hemisphere** | **Criterion** | **Timepoint** | ***beta*** | ***SE beta*** | ***t*** | ***p*** | ***R squared*** | ***Cohen's f2*** |
|---|---|---|---|---|---|---|---|---|---|---|
| **Acquisiton Training** | **Nucleus Accumbens** | **left** | STAI-T | T3 | -0.49 | 0.62 | -0.79 | 0.434 | 0.01 | 0 |
| | | | | T4 | -0.37 | 0.64 | -0.58 | 0.566 | 0.01 | 0 |
| | | | | T5 | -0.71 | 0.59 | -1.20 | 0.236 | 0.02 | 0 |
| | | | BDI | T3 | -0.36 | 0.40 | -0.91 | 0.364 | 0.01 | 0 |
| | | | | **T4** | **-0.09** | **0.47** | **-0.20** | **0.845** | **0.00** | **0** |
| | | | | T5 | 0.18 | 0.36 | 0.51 | 0.609 | 0.00 | 0 |
| | | **right** | STAI-T | T3 | -0.35 | 0.58 | -0.61 | 0.545 | 0.01 | 0 |
| | | | | T4 | -0.57 | 0.62 | -0.91 | 0.364 | 0.01 | 0 |
| | | | | T5 | -0.91 | 0.57 | -1.59 | 0.116 | 0.04 | 0 |
| | | | **BDI** | **T3** | **-0.32** | **0.37** | **-0.86** | **0.393** | **0.01** | **0** |
| | | | | T4 | -0.11 | 0.46 | -0.25 | 0.804 | 0.00 | 0 |
| | | | | T5 | -0.24 | 0.35 | -0.68 | 0.498 | 0.01 | 0 |
| | **Putamen** | **left** | STAI-T | T3 | -0.54 | 0.88 | -0.61 | 0.545 | 0.01 | 0 |
| | | | | T4 | -0.59 | 0.92 | -0.64 | 0.525 | 0.01 | 0 |
| | | | | T5 | -0.10 | 0.86 | -0.12 | 0.907 | 0.00 | 0 |
| | | | BDI | T3 | -0.02 | 0.57 | -0.04 | 0.969 | 0.00 | 0 |
| | | | | T4 | 0.20 | 0.68 | 0.29 | 0.774 | 0.00 | 0 |
| | | | | T5 | 0.40 | 0.51 | 0.79 | 0.434 | 0.01 | 0 |

| Phase | Region | hemisphere | Criterion | Timepoint | *beta* | *SE beta* | *t* | *p* | *R squared* | *Cohen's f2* |
|---|---|---|---|---|---|---|---|---|---|---|
| | **Thalamus** | **left** | STAI-T | T3 | -0.04 | 0.76 | -0.06 | 0.953 | 0.00 | 0 |
| | | | | T4 | 0.26 | 0.82 | 0.32 | 0.751 | 0.00 | 0 |
| | | | | T5 | -0.74 | 0.75 | -0.98 | 0.329 | 0.02 | 0 |
| | | | BDI | T3 | 0.53 | 0.48 | 1.09 | 0.282 | 0.02 | 0 |
| | | | | T4 | -0.03 | 0.60 | -0.05 | 0.959 | 0.00 | 0 |
| | | | | T5 | -0.30 | 0.45 | -0.68 | 0.502 | 0.01 | 0 |
| | **Pallidum** | **right** | STAI-T | T3 | -1.12 | 0.87 | -1.29 | 0.202 | 0.03 | 0 |
| | | | | T4 | -1.05 | 0.91 | -1.15 | 0.253 | 0.02 | 0 |
| | | | | T5 | -2.05 | 0.80 | -2.56 | 0.013 | 0.10 | 0 |
| | | | BDI | T3 | -0.38 | 0.56 | -0.67 | 0.506 | 0.01 | 0 |
| | | | | T4 | -0.81 | 0.67 | -1.21 | 0.232 | 0.02 | 0 |
| | | | | T5 | -0.86 | 0.49 | -1.75 | 0.085 | 0.05 | 0 |
| | **vmPFC** | | STAI-T | T3 | -0.36 | 0.62 | -0.58 | 0.565 | 0.01 | 0 |
| | | | | T4 | -0.41 | 0.65 | -0.62 | 0.537 | 0.01 | 0 |
| | | | | T5 | -0.54 | 0.62 | -0.88 | 0.385 | 0.01 | 0 |
| | | | BDI | T3 | 0.11 | 0.40 | 0.27 | 0.790 | 0.00 | 0 |
| | | | | T4 | -0.18 | 0.48 | -0.37 | 0.714 | 0.00 | 0 |
| | | | | T5 | 0.13 | 0.37 | 0.36 | 0.716 | 0.00 | 0 |
| **Fear Recall** | **Insula** | **left** | STAI-T | T3 | -0.06 | 0.18 | -0.32 | 0.747 | 0.00 | 0 |

| Phase | Region | hemisphere | Criterion | Timepoint | *beta* | *SE beta* | *t* | *p* | *R squared* | *Cohen's f2* |
|---|---|---|---|---|---|---|---|---|---|---|
| | | | | T4 | 0.00 | 0.19 | -0.02 | 0.987 | 0.00 | 0 |
| | | | | T5 | -0.15 | 0.17 | -0.86 | 0.396 | 0.01 | 0 |
| | | | BDI | T3 | -0.03 | 0.12 | -0.26 | 0.798 | 0.00 | 0 |
| | | | | T4 | -0.11 | 0.14 | -0.77 | 0.443 | 0.01 | 0 |
| | | | | T5 | -0.11 | 0.10 | -1.09 | 0.282 | 0.02 | 0 |
| | **Pallidum** | **left** | STAI-T | T3 | 0.09 | 0.18 | 0.47 | 0.641 | 0.00 | 0 |
| | | | | T4 | 0.31 | 0.19 | 1.66 | 0.102 | 0.04 | 0 |
| | | | | T5 | 0.13 | 0.18 | 0.76 | 0.451 | 0.01 | 0 |
| | | | BDI | T3 | 0.06 | 0.12 | 0.53 | 0.595 | 0.00 | 0 |
| | | | | T4 | 0.13 | 0.14 | 0.92 | 0.362 | 0.01 | 0 |
| | | | | T5 | -0.05 | 0.11 | -0.43 | 0.667 | 0.00 | 0 |
| **Reinstatement** | **Amygdala** | **left** | STAI-T | T3 | -0.05 | 0.16 | -0.32 | 0.747 | 0.00 | 0 |
| | | | | T4 | 0.01 | 0.17 | 0.08 | 0.933 | 0.00 | 0 |
| | | | | T5 | 0.02 | 0.16 | 0.11 | 0.916 | 0.00 | 0 |
| | | | BDI | T3 | -0.05 | 0.11 | -0.50 | 0.619 | 0.00 | 0 |
| | | | | T4 | 0.01 | 0.13 | 0.11 | 0.911 | 0.00 | 0 |
| | | | | T5 | -0.06 | 0.09 | -0.68 | 0.501 | 0.01 | 0 |

STAI-T = State-Trait Anxiety Inventory, Spielberger et al., 1983; BDI = Beck Depression Inventory-II, Beck et al., 1996.

#### Comprehensive Rating Results

Supplementary Table 7. Complete results of the mixed-effects models analyzing fear ratings.

| | **Fear_ratings_ACQ** | | | | **Fear_ratings_EXT** | | | | **Fear_ratings_FREC** | | | | **Fear_ratings_RI** | | | |
|---|---|---|---|---|---|---|---|---|---|---|---|---|---|---|---|---|
| *Predictors* | *Estimates* | *CI* | *p* | *df* | *Estimates* | *CI* | *p* | *df* | *Estimates* | *CI* | *p* | *df* | *Estimates* | *CI* | *p* | *df* |
| (Intercept) | 7.02 | 6.07 – 7.98 | **<0.001** | 62.89 | 5.60 | 4.44 – 6.77 | **<0.001** | 66.79 | 9.26 | 8.23 – 10.28 | **<0.001** | 66.44 | 6.91 | 5.90 – 7.92 | **<0.001** | 67.87 |
| CS type1 | 1.83 | 0.95 – 2.72 | **<0.001** | 185.18 | 2.09 | 1.32 – 2.85 | **<0.001** | 193.54 | 5.75 | 4.95 – 6.54 | **<0.001** | 186.44 | 1.48 | 0.67 – 2.29 | **<0.001** | 182.52 |
| Recent adversity [1] | 0.56 | -1.06 – 2.18 | 0.490 | 65.04 | -0.12 | -2.10 – 1.85 | 0.901 | 66.70 | 0.39 | -1.35 – 2.14 | 0.654 | 66.83 | -1.38 | -3.08 – 0.31 | 0.107 | 65.69 |
| Fear ratings T0 | 0.78 | 0.68 – 0.88 | **<0.001** | 183.16 | 0.68 | 0.56 – 0.81 | **<0.001** | 193.04 | 0.36 | 0.26 – 0.46 | **<0.001** | 186.67 | 0.47 | 0.32 – 0.63 | **<0.001** | 178.47 |
| CS type1 × Recent adversity [1] | -0.48 | -1.87 – 0.90 | 0.493 | 188.00 | 0.05 | -1.11 – 1.21 | 0.928 | 195.33 | -0.59 | -1.60 – 0.42 | 0.248 | 187.08 | 0.33 | -0.93 – 1.59 | 0.602 | 178.59 |
| Time1 | | | | | | | | | 0.47 | -0.15 – 1.10 | 0.139 | 191.31 | -2.28 | -3.17 – -1.38 | **<0.001** | 181.13 |
| Time1 × CS type1 | | | | | | | | | 0.87 | 0.27 – 1.47 | **0.005** | 186.06 | -0.56 | -1.32 – 0.19 | 0.143 | 180.71 |
| Time1 × Recent adversity [1] | | | | | | | | | -0.30 | -1.31 – 0.71 | 0.555 | 190.73 | 0.76 | -0.50 – 2.02 | 0.234 | 176.44 |
| (Time1 × CS type1) × Recent adversity [1] | | | | | | | | | -0.08 | -1.08 – 0.92 | 0.876 | 186.80 | 0.23 | -1.03 – 1.50 | 0.715 | 178.42 |

| | | | | |
|---|---|---|---|---|
| **Random Effects** | | | | |
| $\sigma^2$ | 27.72 | 20.45 | 15.11 | 22.37 |
| $\tau_{00}$ | 2.54 $_{Subject}$ | 9.88 $_{Subject}$ | 7.89 $_{Subject}$ | 4.94 $_{Subject}$ |
| ICC | 0.08 | 0.33 | 0.34 | 0.18 |
| N | 67 $_{Subject}$ | 69 $_{Subject}$ | 69 $_{Subject}$ | 69 $_{Subject}$ |
| Observations | 250 | 264 | 261 | 246 |
| Marginal $R^2$ / Conditional $R^2$ | 0.612 / 0.644 | 0.446 / 0.626 | 0.735 / 0.826 | 0.444 / 0.545 |

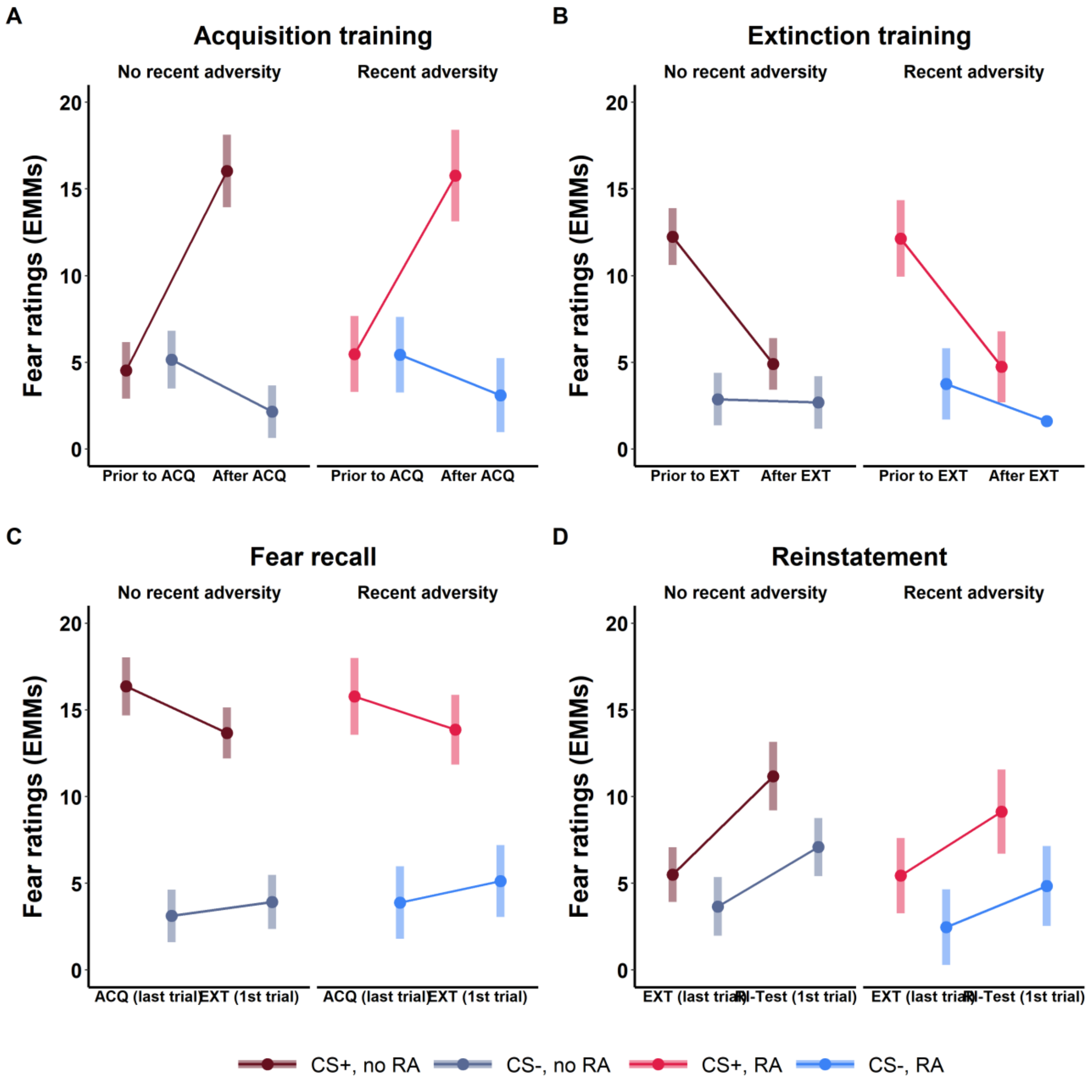


Supplementary Figure 4: Estimated marginal means (EMMs) of fear ratings at T1 (controlled for T0 responding) during acquisition training (A), extinction training (B), for fear recall (C) and reinstatement (D) separated for CS type and exposure to recent adversity. CS = conditioned stimulus. ACQ = acquisition training, EXT = extinction training, RI = reinstatement.

72. Scharfenort R, Menz M, Lonsdorf TB. Adversity-induced relapse of fear: Neural mechanisms and implications for relapse prevention from a study on experimentally

induced return-of-fear following fear conditioning and extinction. Transl Psychiatry. 2016;6:e858. DOI: 10.1038/tp.2016.126
